# A tradeoff between robustness to environmental fluctuations and speed of evolution

**DOI:** 10.1101/834234

**Authors:** Max Schmid, Maria Paniw, Maarten Postuma, Arpat Ozgul, Frédéric Guillaume

**Author notes:** **statement of authorship:** MS and FG conceived the study, and all authors further developed the approach. MS and FG derived the mathematical equations. MS performed the simulations, and analyzed output data. MS and FG wrote the paper. All authors substantially contributed to revisions of early drafts of this work. **data accessibility statement** The authors hereby confirm that the simulation data supporting the results will be archived in an appropriate public repository and the data DOI will be included at the end of the article.

## Abstract

Organisms must cope with both short- and long-term environmental changes to persist. In this study we investigated whether life histories trade-off between their robustness to short-term environmental perturbations and their ability to evolve directional trait changes. We could confirm the tradeoff by modeling the eco-evolutionary dynamics of life-histories along the fast-slow pace-of-life continuum. Offspring dormancy and high adult survival rates allowed for large population sizes to be maintained in face of interannual environmental fluctuations but limited the speed of trait evolution with ongoing environmental change. In contrast, precocious offspring maturation and short-living adults promoted evolvability while lowering demographic robustness. This tradeoff had immediate consequences on extinction dynamics in variable environments. High evolvability allowed short-lived species to cope with long-lasting gradual environmental change, but came at the expense of more pronounced population declines and extinction rates from environmental variability. Higher robustness of slow life-histories helped them persist better on short timescales.

## Introduction

Environmental conditions are rarely constant but fluctuate between years and average conditions change gradually over time, and both dynamics pose a considerable challenge for species (Bürger and Lynch, 1995; Boyce et al., 2006; Van De Pol et al., 2010; Vázquez et al., 2015). For instance average global temperatures change successively in the course of long-term glacial cycles or during ongoing anthropogenic climate change, but also vary on shorter time scales, for instance with variations between years (Stocker et al., 2013; Ummenhofer and Meehl, 2017). Given that the environmental tolerance of all species is finite, the exposure to short- or long-term environmental changes can result in higher mortality, lower recruitment, and population declines. While a dynamic environment thus demands species to constantly adapt and adjust, several adaptive strategies exist to do so (Parmesan, 2006; Aitken et al., 2008; Bell and Gonzalez, 2009; Doak and Morris, 2010; Duputié et al., 2015; Cayuela et al., 2017; McDonald et al., 2017). Life-history characteristics affect both the robustness of a population against short-term environmental fluctuations (Caswell, 2001; Boyce et al., 2006; Morris et al., 2008; Tuljapurkar et al., 2009), and the speed of adaptive evolution in response to gradual environmental change (Engen et al., 2011; Barfield et al., 2011; Orive et al., 2017). However, life histories’ ability of to cope with one kind of environmental variability might come at a cost to cope with another kind of dynamic.

Stochasticity in vital rates (survival and fecundity) from random environmental fluctuations reduces the long-run fitness of populations and may depress population sizes, thereby increasing extinction risks (Tuljapurkar, 1982; Lande and Orzack, 1988; Tuljapurkar et al., 2003; Engen et al., 2005; Sæther et al., 2013). How severely environmental variability feeds back on population growth then depends on the species’ life cycle, especially on its position along the fast-slow continuum and its reproductive strategy. Theory shows that species with longer generation times are more robust to fluctuations than short lived species, as are more iteroparous compared to semelparous species (Tuljapurkar, 1990; Tuljapurkar et al., 2009; Salguero-Gómez et al., 2017). Empirically, Morris et al. (2008) found a relation between the life expectancy of 36 plant and animal species and their sensitivity to environmentally driven variation in vital rates. Life histories with long generation times experienced less pronounced demographic consequences from stochasticity in vital rates than short-lived species (see also Dalgleish et al., 2010; Sæther et al., 2013). A life history’s robustness against environmental fluctuations further depends on the proportion of individuals exposed to environmental extremes. For instance, long-lived plants often exhibit highly tolerant age or stage classes, like seeds or adult trees, that can overcome detrimental conditions much better than seedlings or juveniles (Petit and Hampe, 2006; Buoro and Carlson, 2014) and help to “disperse through time” (Evans and Dennehy, 2005; Buoro and Carlson, 2014).

Life-history strategies also shape the rate of trait evolution when the environment changes gradually (Lande, 1982; Charlesworth, 1994; Ellner and Hairston, 1994; Engen et al., 2009, 2011; Barfield et al., 2011; Orive et al., 2017; Cotto et al., 2019). The generation time of a species, which affects the pace of life, is an indicator for how fast selection can translate into changes of the gene pool. The lower the generation time, the greater is the evolutionary response to environmental change per unit of time (Lande, 1982; Vander Wal et al., 2013; Carlson et al., 2014; Orive et al., 2017). With life stages highly tolerant to environmental variation, life-history strategies further control the proportion of individuals exposed to the environment. Those genotypes “hidden” in the seedbank or in highly tolerant adult stages are not under selection and thereby delay genetic changes at the population level (Templeton and Levin, 1979; Hairston and De Stasio Jr., 1988; Barfield et al., 2011; Orive et al., 2017). Together with the life-history effects on robustness mentioned above, we can expect that short-lived species will have faster evolutionary trait responses but will suffer more from a higher demographic sensitivity. In contrast, life histories with longer generation times should be demographically more robust but less evolutionarily responsive to changes in their environments. This tradeoff between demographic robustness and evolvability is in fact implicit from previous theoretical work. Lande (1982) and Barfield et al. (2011) described the evolutionary response to selection as a function of the growth rate sensitivity to vital rate changes, which is a quantity related to robustness against short-term fluctuations. The tradeoff was also suggested by the results of Templeton and Levin (1979), Hairston and De Stasio Jr. (1988), and Orive et al. (2017), who found that the presence of a seed bank and elevated iteroparity slowed down the rate of phenotypic trait evolution.

While several findings point towards a robustness-evolvability tradeoff for life-history strategies, other studies hint at potential mechanisms to overcome this tradeoff. Life histories with overlapping generations were suggested to harbour higher levels of additive genetic variance in fluctuating environments than life histories with non-overlapping generations, a process known as the genetic storage effect (Ellner and Hairston, 1994; Sasaki and Ellner, 1997; Svardal et al., 2015). Given that the likelihood of a successful evolutionary rescue increases with the extent of standing genetic variation (Carlson et al., 2014), life histories with highly robust life stages might be able to accumulate higher levels of genetic variance, and thus also evolve faster (Yamamichi et al., 2019). All in all, this shows that life-history strategies can have large effects on evolutionary and demographic dynamics, and on the success of evolutionary rescue. However, the exact resolution of the tradeoff between evolvability and demographic robustness is not solved yet and will likely depend on the details of the species’ life cycle and on the type of environmental variation.

Here, we first characterized the tradeoff between demographic robustness and evolvability for four life-history strategies spanning the fast-slow continuum and different levels of iteroparity in stage-structured populations. Our analytical analysis of the tradeoff is based on classical matrix population modeling (Caswell, 2001) and on the quantitative genetics formalism developed by Lande (1982) and Barfield et al. (2011). We show that a linear tradeoff exists between sensitivities of the equilibrium population size and growth rate, and the rate of evolution of a quantitative trait. In a second step, we used individual-based simulations to model the eco-evolutionary dynamics of a single population for the same life histories subject to both random inter-annual fluctuations and long-term gradual changes of the optimum trait value. The simulations further allowed us to account for demographic and genetic stochasticity and evolving additive genetic variance. We could thus study the effect of the tradeoff between robustness and evolvability on the population extinction risk from environmental changes.

## Models

We modeled a hermaphroditic species with three life stages (offspring (*n*_1_), juveniles (*n*_2_), and adults (*n*_3_)) in a single random mating population. Environmental fluctuations only affected juvenile survival via stabilizing selection on a quantitative trait. This corresponds to situations where the focal trait is only expressed at the juvenile stage (e.g., growth), and where offspring and adult individuals have much higher environmental tolerance (e.g., frost or drought tolerance), or dwell in much more stable habitats (e.g., Petit and Hampe, 2006; Van De Pol et al., 2010; Marshall et al., 2016).

We focused on four distinct life-history strategies along two axes, timing of maturation and degree of iteroparity, with two cases each: precocious (**pre**) and delayed (**del**) maturation, iteroparity (**ite**) and semelparity (**sem**). The life histories with combinations of delayed maturation and iteroparity (*del-sem, del-ite, pre-ite*) had three stages and overlapping generations (Fig. 1a). The precocious, semelparous life cycle (*pre-sem*) had non-overlapping generations modeled with only two stages, juveniles (offspring maturing immediately) and adults reproducing once and dying (Fig. 1b).

**Figure 1:**
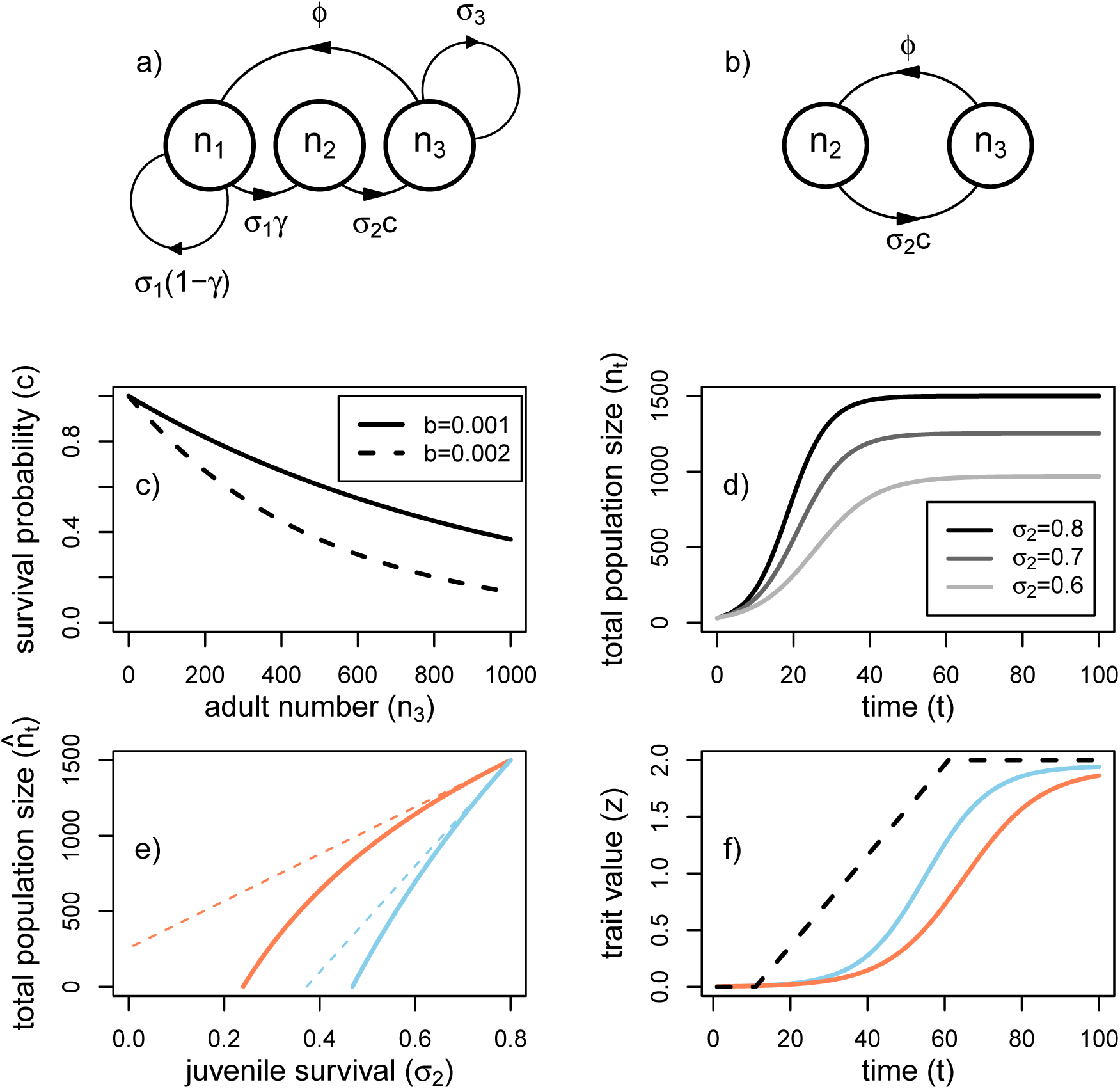
The life-history plot in a) illustrates the three-stage life cycle with delayed maturation and long-living adults (*del-sem, del-ite, pre-ite*), while b) shows the biennial two-stage life cycle (*pre-sem*), together with the relevant vital rates. Graph c) demonstrates the effect of negative density dependence on juvenile survival in the three-stage life cycle based on the Ricker function. Graph d) illustrates the temporal change of total population size over time following logistic growth such that an equilibrium mean population size is achieved from negative density dependence. By reducing the average juvenile survival 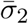, for instance from environmental perturbations, the population size at equilibrium 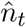 is reduced. We used the sensitivity of the equilibrium population size to juvenile survival 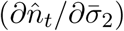 as a measure for environmental robustness. In graph e), two life histories are shown with the same equilibrium population size (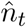, the two solid lines —) when environmental fluctuations are absent (at 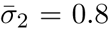) but with different sensitivities to reductions in juvenile survival from interannual fluctuations. The slope of the dotted lines (…) in graph e) represents 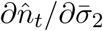 as how quickly 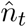 responds to small reductions in 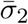. Graph f) illustrates the scenario with directional environmental change and the the corresponding trait evolution of two life histories (the two solid lines —). Environmental change is realized via directional changes of the phenotypic optima (*θ*, dashed line −−).

We modeled two growth scenarios, one of an exponentially growing population, and one of a population that reached an equilibrium population density 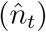 from intra-specific competition (Fig. 1c,d).

### Life-history parameters

Offspring, juvenile and adult individuals survived to the next year with average probability 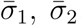, and 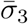, respectively (Fig. 1a,b). Offspring exclusively resulted from the sexual reproduction of hermaphroditic adults with an average per capita fecundity of 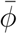. Offspring matured to juveniles with mean maturation probability 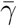 and stayed in the offspring stage with probability 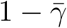.

With density regulation, average survival of juveniles was reduced by intra-specific competition 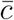 following the Ricker function 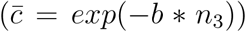 and thus declined with increasing adult number *n*_3_ for overlapping generations, and with juvenile number *n*_2_ for non-overlapping generations, depending on the competition coefficient *b*. We also modeled a Beverton-Holt density regulation and present the results in the Supplementary Material. In absence of density regulation, competition survival was a constant and set to 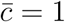.

The demographic dynamics of the three-stage life cycles *del-sem, del-ite*, and *pre-ite* were modeled by the following matrix population model (MPM):

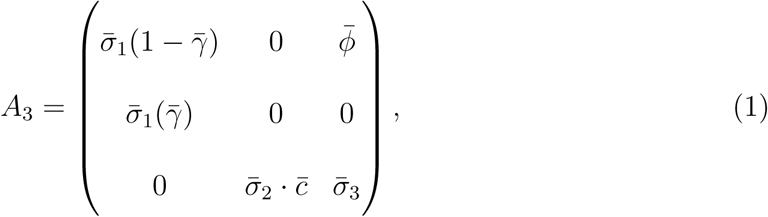

and the two-stage life cycle *pre-sem* was represented by the following MPM:

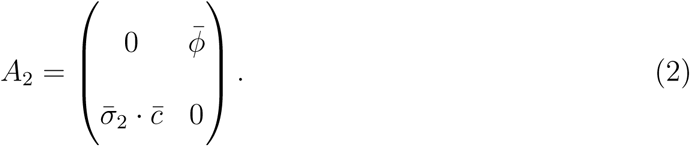

### Population sensitivity to environmental fluctuations

For exponentially growing populations, the robustness of a life history against environmental fluctuations can be seen as the sensitivity of the average population growth rate 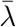 to average juvenile survival 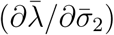. The growth rate sensitivity 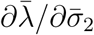 relates to demographic robustness as it determines by how much variation in vital rates translate into variation of the growth rate, and thereby by how much environmental variability reduces the long run growth rate of a population (see Tuljapurkar, 1982; Lande, 2007). We computed 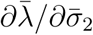 for each life-history variant using explicit derivations obtained with Mathematica 12.1 (Wolfram Research, 2020).

With density regulation, an alternative measure for demographic robustness is the sensitivity of the total population size at equilibrium 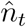 to juvenile survival (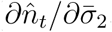, see also Fig. 1e). The sensitivity 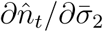 determines by how much a reduction in juvenile survival from environmental perturbation feeds back on population density. As 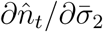 is not necessarily proportional to 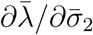 (Takada and Nakajima, 1998; Grant and Benton, 2000, 2003; Caswell et al., 2004), we present the results for both sensitivity measures in parallel. We derived the partial derivative of 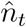 with respect to 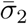 depending on single vital rates as detailed in Appendix A and B.

### The rate of trait evolution

We quantified the speed of evolution for a life history as the average asymptotic trait change 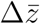 per time step *t* from genetic evolution (not plasticity) in response to directional selection. We will thus use 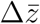 as a measure of evolvability *e* as defined by Hansen et al. (2019): 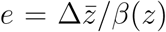, or 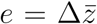 for standardized *β*(*z*) = 1. The speed of evolution for quantitative traits in a stage-structured population was derived by Barfield et al. (2011):

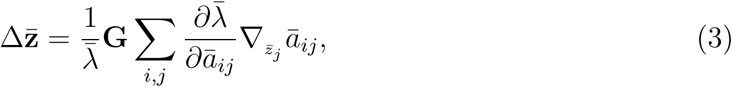

where 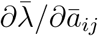 is the sensitivity of the average growth rate 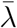 to changes in the average vital rates *ā*_*ij*_ (Caswell, 2001), and 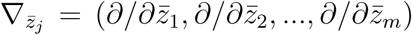 is the gradient operator with respect to trait means at stage *j* (Barfield et al., 2011). Equation 3 was obtained under the assumption of density-independent population dynamics leading to a stable stage distribution (SSD) and a transition matrix **A** that is nearly constant over time, weak selection, and same mean phenotype 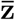 and genetic variance **G** for all stages *j* (see discussion in Barfield et al., 2011, Appendix B). Importantly, equation 3 shows that the rate of evolution depends linearly on the growth rate sensitivity to variation in vital rates and on the sensitivity of vital rates to change in average trait values. Equation 3 also accounts for the changes of vital rates caused by changes in trait value *z*. We consider here that variation in survival is caused by the phenotypic mismatch of the population with its environment, which only affects the vital rate of juveniles *ā*_32_. The relationship between phenotypic trait value *z* and *ā*_32_ is given by 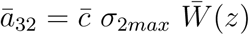, the product of maximum juvenile survival (*σ*_2*max*_), competition survival 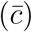, and average juvenile survival depending on the mismatch between individual phenotypes and the environment 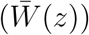.

For a species with three life-history stages with one trait under selection in juveniles (*z*_2_), the asymptotic rate of evolution can be derived from Equation 3 as detailed in Appendix A:

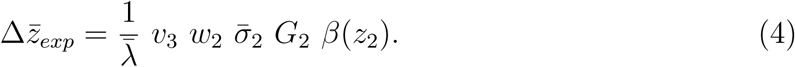

The asymptotic rate of trait change 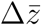 thus varies with the additive genetic variance in juveniles (*G*_2_), the selection gradient on a single juvenile trait (*β*(*z*_2_)), the fraction of individuals exposed to selection (*w*_2_, the proportion of juveniles at SSD), the survival probability of these individuals 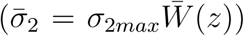, and their contribution to the subsequent generations (*v*_3_, the reproductive value of adults standardized such that *v*^*T*^ *w* = 1) (equation 8, Barfield et al., 2011). The rate of evolution 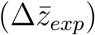 can thus be expressed as a function of the vital rates by deriving expressions for *w*_2_ and *v*_3_ (see Appendix A, and Appendix B for the 2-stage life cycle).

Beside the derivations for exponentially growing populations, we also derived evolvability for populations at equilibrium population density (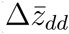, see Appendices A and B).

#### Standardization of the life cycles

We standardized life histories for uniform maximum population sizes 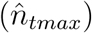 in absence of environmental variation and for uniform maximum population growth rates (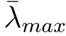, the intrinsic population growth rate in absence of density regulation when 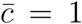). Standardization to equal 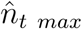 was necessary because 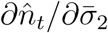 is a function of 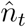, and 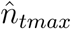 determines how quickly population size will reach a threshold under which demographic stochasticity will lead to extinction. Standardization to equal 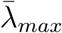 allows us to compare populations with similar growth under non-equilibrium conditions. With this standardization we could compare our four life histories while keeping key demographic properties constant. Using a numeric approach, we adjusted the fecundity 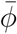 to reach equal 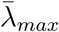 across life histories, and a standardized form of the competition coefficient *b* to reach equal 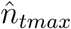 (see Appendix A and B).

Using the derivations from Appendix A and B, we explored the effect of the vital rates 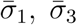, and 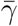 on the tradeoff between demographic robustness 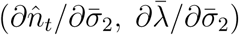 and evolvability (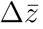, with *G* = *β*(*z*_2_) = 1).

#### Individual-based simulations

We ran individual-based simulations to test our mathematical derivations and account for genetic and demographic stochasticity, and an evolving additive genetic variance. Simulations were run with Nemo (Guillaume and Rougemont, 2006) extended for stage-structured populations (Cotto et al., 2017, 2020). We simulated a single panmictic population of hermaphroditic individuals. The population reached an equilibrium population size by density-dependent juvenile survival. A single quantitative trait was under hard selection in the juvenile stage. The following life cycle events occurred consecutively each year: density-dependent regulation (of juvenile survival using the Ricker function), viability selection (removal of juveniles according to their survival probability which varied with the distance between trait value and environment), sexual reproduction (independent from the environmental state or population density), and stage transitions according to the MPM.

#### Life-history parameters

We modeled the four generic life-history strategies presented in the previous section. We standardized all four life-history strategies for the same total population size in absence of environmental fluctuations (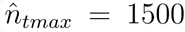 by adjusting *b*) and the same maximum population growth rate (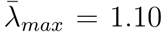 or 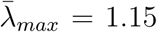, by adjusting 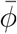). The single vital rates are listed in table S1. With these simulation scenarios we tried to span a range of key life-history characteristics by accounting for differences in “pace of life” along the slow-fast continuum and for differences in the reproductive strategy (see Fig. S15, and Stearns, 1983; Gaillard et al., 1989, 2005; Salguero-Gómez, 2017).

#### Genetic setup and selection model

We simulated the evolutionary dynamics of adaptation to a single environmental condition via the evolution of a single quantitative trait (*z*) under stabilizing selection in the juvenile stage using the classical Gaussian survival function *W* (*z*) = *exp*(−(*z* − *θ*)^2^*/*2*ω*^2^), with *θ* the trait phenotypic optimum, and *ω*^2^ the width of the survival function. We set *ω*^2^ = 4 (corresponding to stronger selection) or *ω*^2^ = 16 (representing weaker selection). Each individual carried *l* = 20 unlinked loci, contributing additively to the trait *z*. We used a continuum-of-allele model where mutational effects, picked from a Gaussian distribution centered on zero with mutational variance *α* = 0.1, were added to the existing allelic values. The mutation rate was *µ* = 0.0001 so that the mutational phenotypic variance was *V*_*M*_ = 2*lµα* = 0.001. We excluded random environmental effects on phenotype expression (e.g., developmental noise) for simplicity.

#### Environmental scenarios

We simulated environmental variation between years together with gradual changes of the average condition by perturbing the phenotypic optimum *θ* (e.g., see Chevin et al., 2017). Each year, the position of the phenotypic optimum for the entire population was picked from a Gaussian distribution 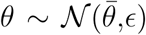 with the average environment 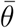 and variance *ϵ*. We thus simulated environmental fluctuations between years that were independently and identically distributed (e.g., Caswell, 2001).

To allow the population genetic variance to reach mutation-selection-drift balance, we ran burn-in simulations for 50’000 years for each life history. Burn-in simulations were run with a constant average environment of 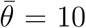 for six different degrees of environmental fluctuations (*ϵ* = 0.0, 0.2, 0.5, 1.0, 1.5, 2.0).

To asses the population size sensitivity of life-history strategies to interannual environmental stochasticity, we recorded population demographic characteristic (e.g., population size 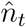) every 10 years for an additional period of 500 years. A life history’s population size sensitivity was assessed as the steepness of declining average population sizes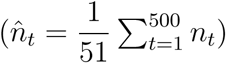 with increasing degrees of interannual environmental fluctuations *ϵ*.

To asses the speed of trait evolution, we then initiated directional environmental change of the average environmental condition 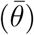 after burn-in, while still simulating interannual variation on top of the changing mean. The average phenotypic optimum started to change at *t*=100 with a rate of 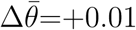, for 200 and 800 years. We quantified the speed of evolution as the time to reach 90% of the novel phenotypic optimum (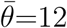, or 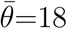). We did not use the maximum rate of phenotypic change per time step because all life histories either went extinct or adopted trait changes as fast as the rate of environmental change 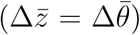.

## Results

### Mathematical models

We found a tradeoff between life-history effects on the rate of evolution and their robustness against environmental perturbations, at least when comparing life histories with the same maximum growth rate. The tradeoff is explicit in the positive, linear relationship between the rate of evolution 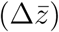 and the sensitivity of the population size to juvenile survival (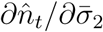 and 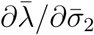) for both the three-stage and the two-stage model (Fig. 2). Changes in vital rates that enhanced the rate of evolution also increased the sensitivity to environmental perturbations, and vice versa.

**Figure 2:**
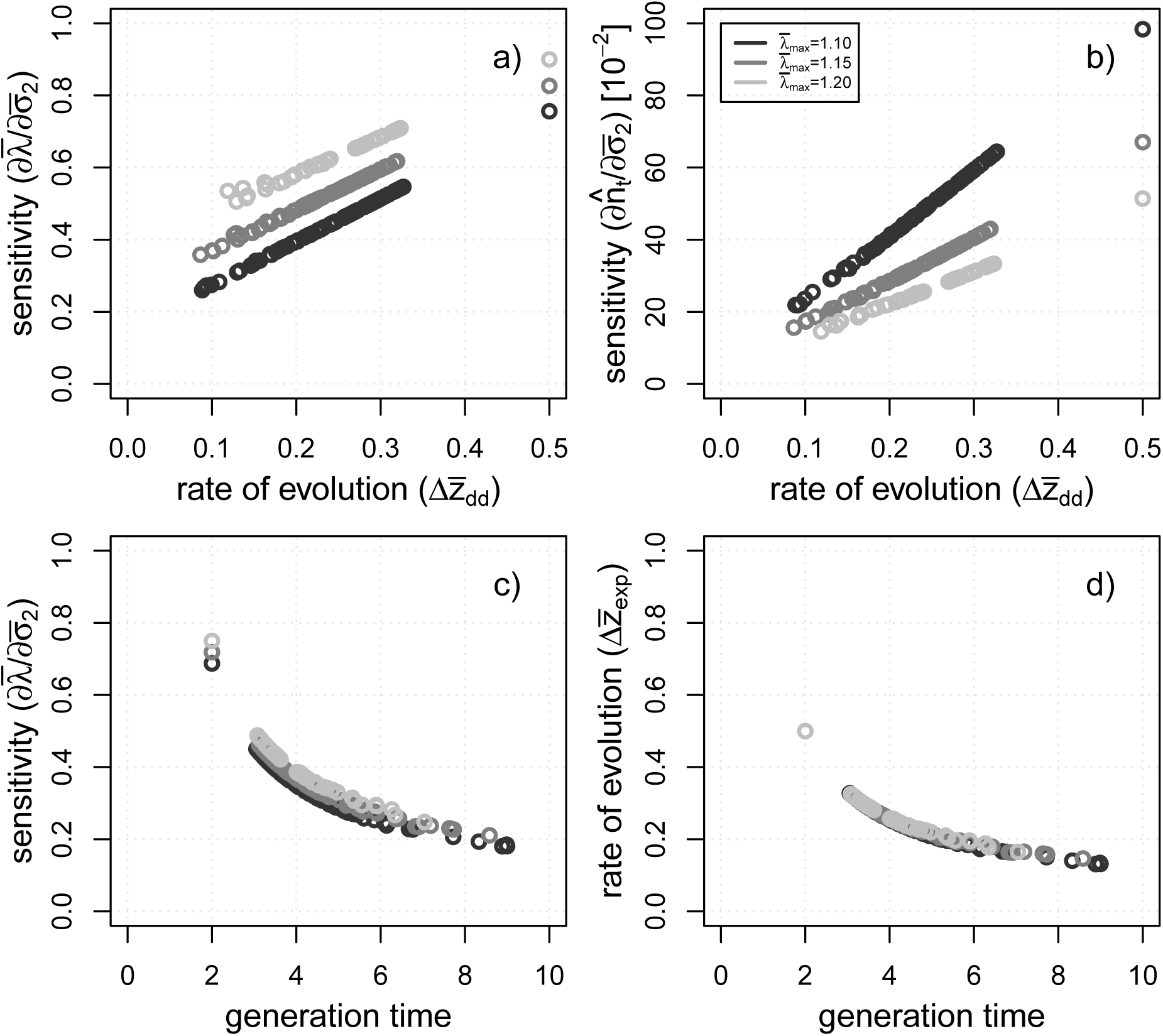
A life history’s tradeoff between evolvability 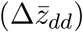 and robustness 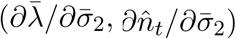 with density regulation at 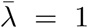 are illustrated in a) and b). In addition, the relation between robustness, evolvability, and generation time (all evaluated at exponential growth) are shown in c) and d). Each circle illustrates a random combination of vital rates (with 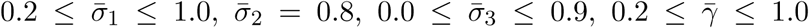). Fecundity 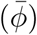 was recalculated for each vital rate combination to standardize the maximum population growth rate 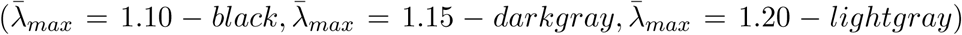. For a) and b) the strength of competition (*b*) was re-calculated for each life-history variant to standardize for the same total population size (*n*_*t*_=1’500) respectively. We then computed the rate of evolution (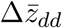 with *G*_2_*β*(*z*_2_) = 1), the population size sensitivity 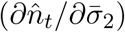, and the growth rate sensitivity 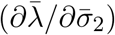. Generation time was computed with the *popbio* package (version 2.7, Stubben and Milligan, 2007) in R (version 3.6.2, R Core Team, 2019). The three single points on the right (respectively on the left) of the plots were derived for the 2-stage life cycle (with 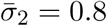), while the rest of the points illustrate parameter combinations for 3-stage life cycles.

A higher degree of iteroparity (high adult survival 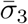 Fig. 3a,d), delayed maturation (low maturation rate 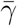; Fig. 3c,f), and thus longer generations times (Fig. 2c,d) increased the robustness of the population size to environmental fluctuations, but reduced the rate of trait evolution. In contrast, precocious maturation and semelparous adults allowed for fast evolution but reduced the robustness to environmental variability. We obtained qualitatively similar results for the sensitivity of the growth rate 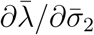, at least for a given maximal growth rate (Fig. 3).

**Figure 3:**
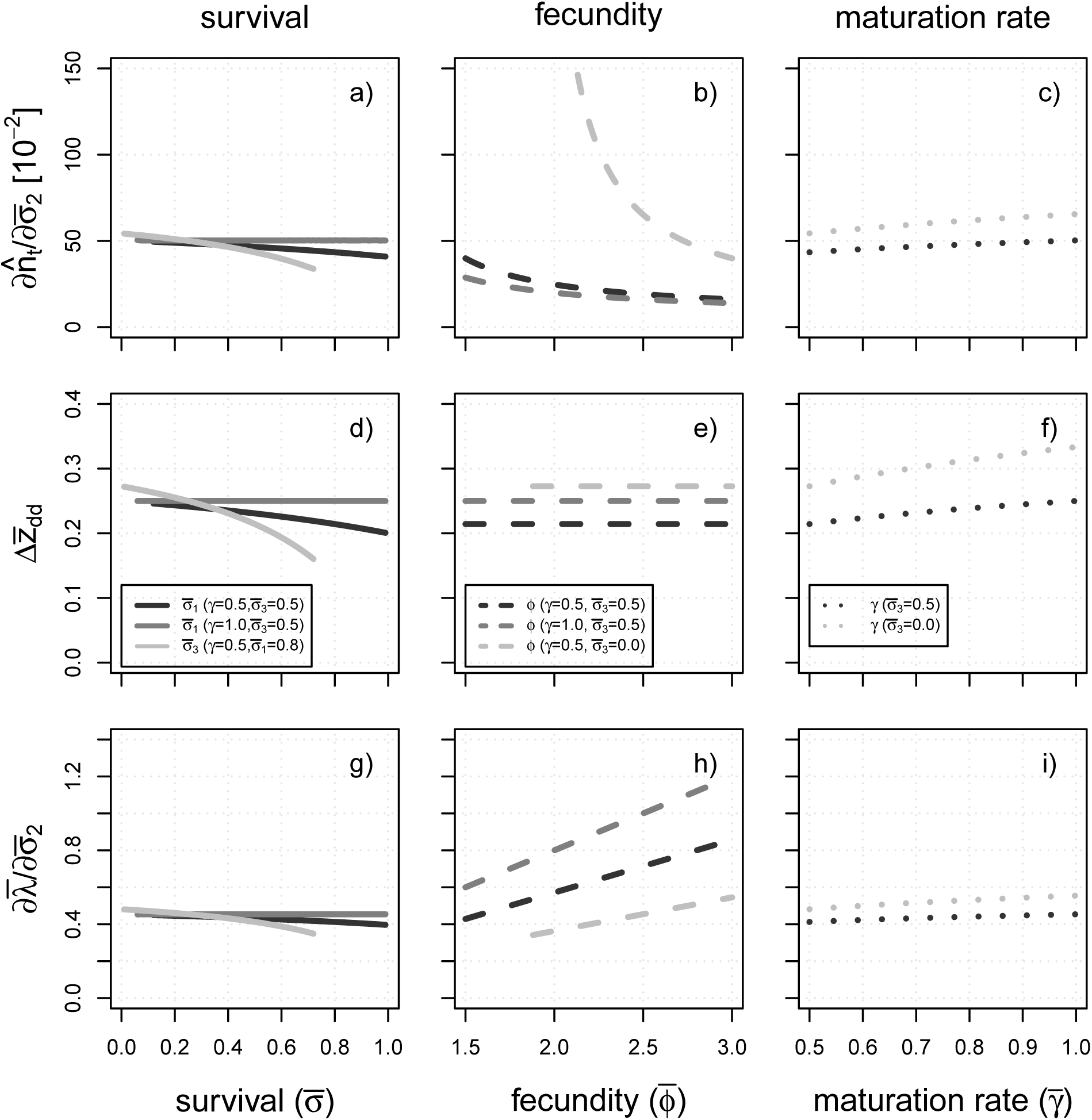
The graphs illustrate the analytical results for the sensitivity of the total population size to juvenile survival (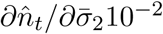, a-c), the rate of evolution (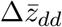, d-f), and the sensitivity of the population growth rate (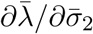, g-i) for three-stage life cycle depending on survival rate 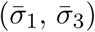, fecundity 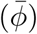, and maturation probability 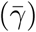. The sensitivity measures 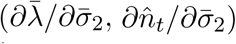 were estimated at equilibrium population size in absence of interannual environmental variation at 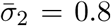. The maximum growth rate has been standardized for 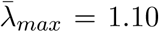 by fecundity adjustments, except for the case with varying fecundity (b,e,h). Each parameter combination has been standardized for the same equilibrium population size in absence of inter-annual environmental fluctuations 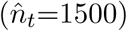 via adjustments of the strength of competition (*b*). If not specified otherwise, the following parameter values have been used : 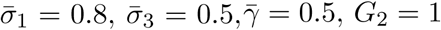, and *β*(*z*_2_) = 1.

The maximum rate of population growth 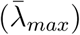 shaped both the population size and growth rate sensitivity to environmental fluctuations, even though it had no effect on the rate of evolution at carrying capacity 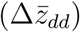, and only minor effects on evolvability

with exponential growth 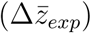. Higher 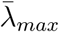 reduced 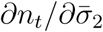 and thus helped to maintain higher population densities with short-term environmental perturbations (Fig. 2b). However, higher 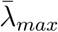 increased 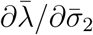 (Fig. 2a, Fig. 3b,h).

Results for a Beverton-Holt instead of a Ricker function for density regulation were qualitatively similar. The linear robustness-evolvability tradeoff remained (Fig. S6). Only the population size sensitivity 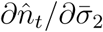 was higher with a Beverton-Holt regulation function (Fig. S7).

### Individual-based simulations

We could confirm the tradeoff between evolvability and robustness also in our individual-based simulations as the genetic storage effect was not strong enough to overcome the tradeoff. Life-history strategies with high robustness had the slowest rates of evolution, and vice versa. The average population size maintained at constant average conditions during burn-in declined with increasing environmental fluctuations for all life cycles below 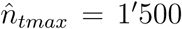, while life-history strategies differed from each other in the extent of 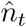 reductions (Fig. 4b, Fig. 5a). The onset of gradual environmental change further reduced juvenile survival and evoked a decline in juvenile number and total population size (Fig. 4b,d), before trait evolution allowed to adapt to the changing conditions (Fig. 4a) and recover in a later phase of environmental change (Fig. 4d). Evolutionary rescue resulted in a u-shaped curve in total population size *n*_*t*_, with a population size recovery already during environmental change or only after environmental change has stopped (Fig. 4b).

**Figure 4:**
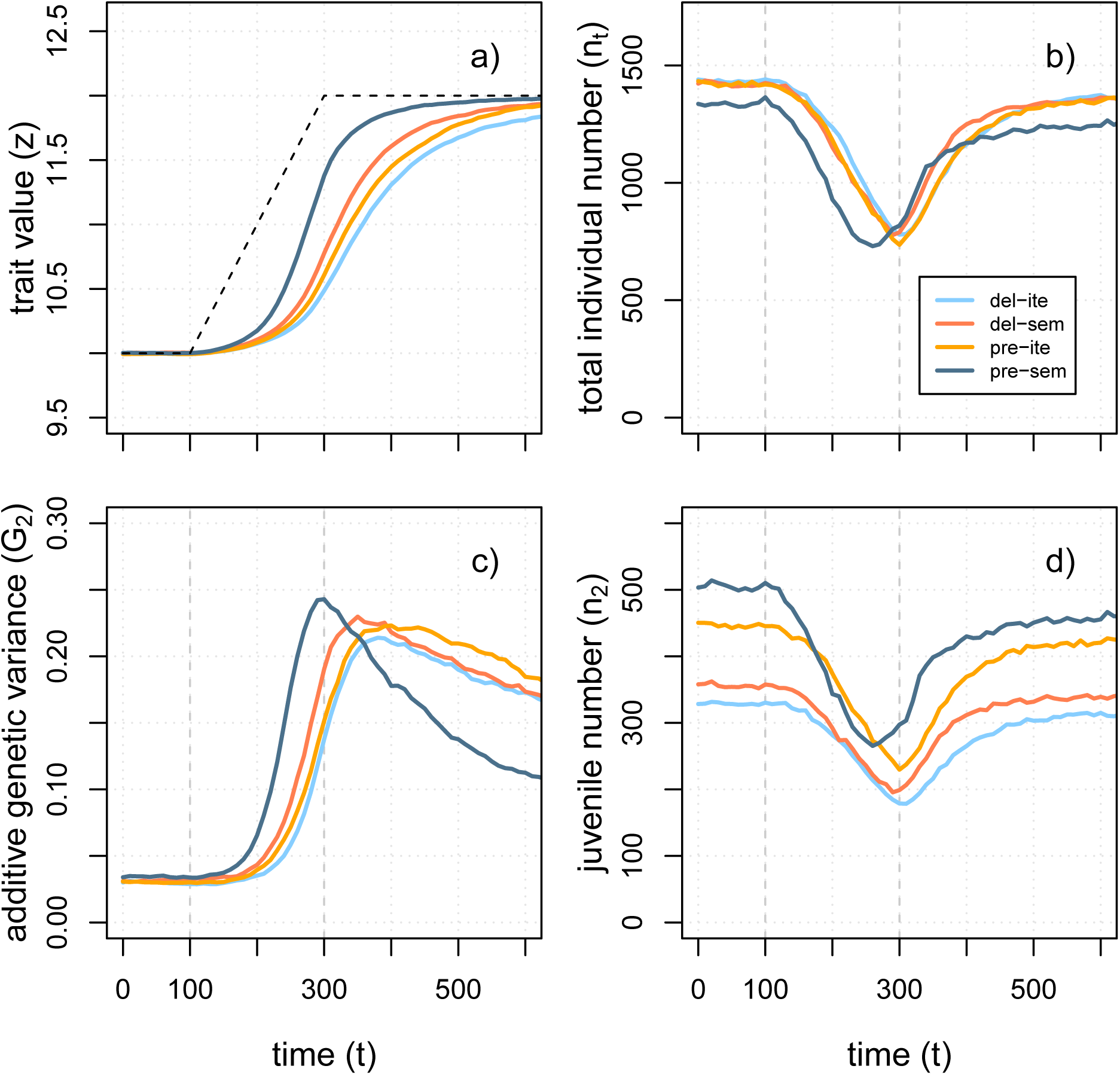
The graphs show the results of individual-based simulations for the mean trait value *z* (a), the total population size *n*_*t*_ (b), the additive genetic variance *G*_2_ (c), and the juvenile number *n*_2_ (d) for the four life histories (*del-sem, del-ite, pre-ite, pre-sem*) over 600 years. Environmental change started at t=100 and was realized by a gradual shift in the phenotypic optimum (−− black dashed lines) from *θ* = 10 to *θ* = 12 by t=300 (a). The between-year environmental fluctuations were set to *ϵ* = 0.2 and life histories were standardized for 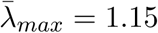. Average population sizes (*n*_*t*_) were standardized to *n*_*t*_ = 1500 in absence of interannual environmental fluctuations, and were depressed as result of between-year fluctuations and directional environmental change as result of a fitness reduction in juveniles (a). All values of the total population size and trait values represent the arithmetic means over all of the 100 replicates that did not go extinct.

**Figure 5:**
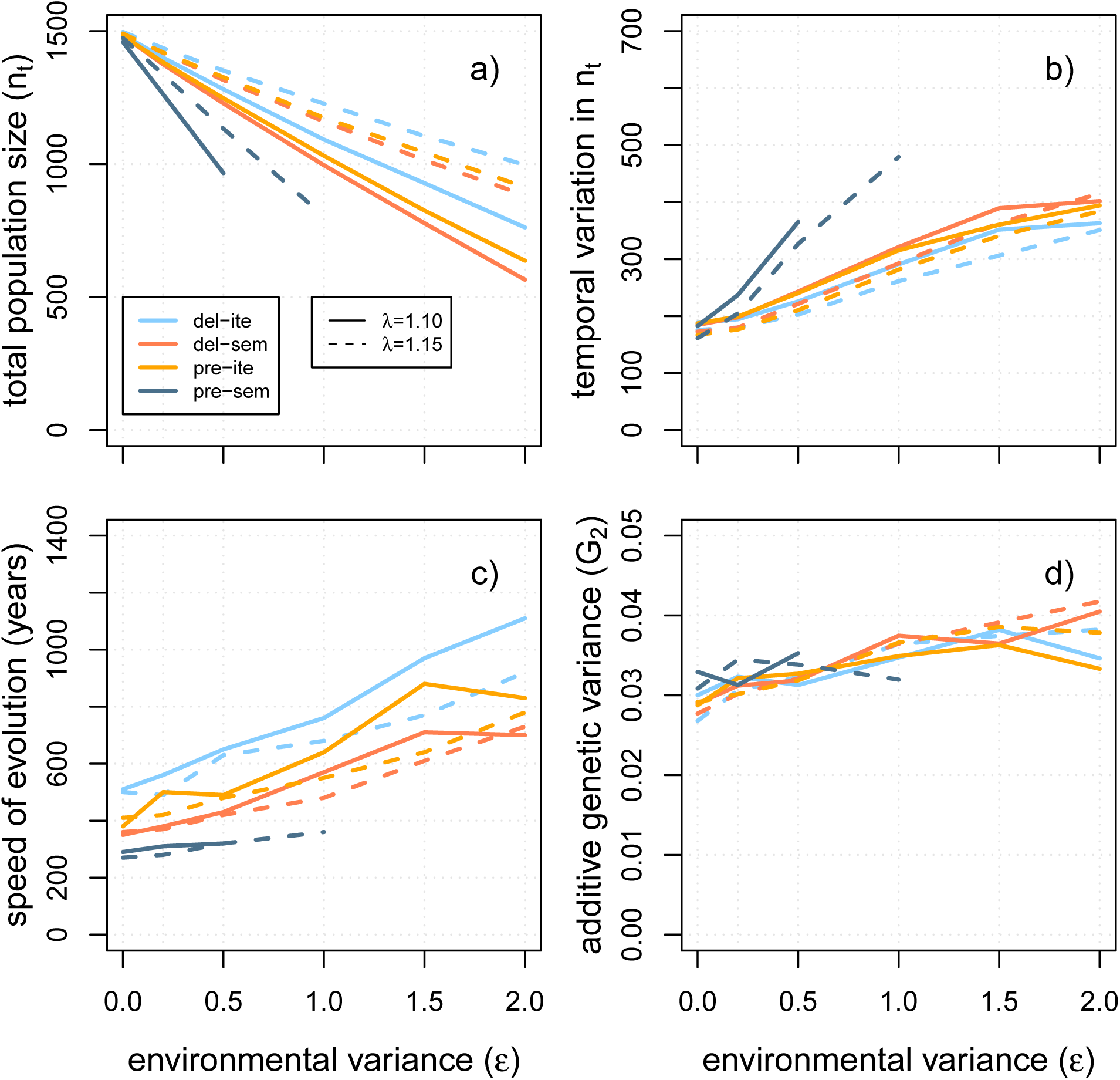
The graphs illustrate the total population size *n*_*t*_ (a), the variability in *n*_*t*_ over time (90 % quantile of *n*_*t*_ over 500 years, b), the speed of evolution (the average number of years to reach the trait value *z* = 11.8, c), and the additive genetic variance (*G*_2_, d) are shown depending on the interannual environmental variability (*ϵ*). The simulation results are shown for four life histories (*del-sem, del-ite, pre-ite, pre-sem*) that have been standardized for two maximal population growth rates 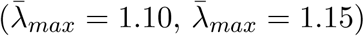 with environmental change lasting for 200 years.

#### Population size sensitivity to environmental variability

The sensitivity of the total population size 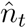 to juvenile survival (Fig. 5a,b) matched with our analytical predictions (Tab. S1). The biennial, two-stage life cycle (*pre-sem*) had the greatest sensitivity, shown in the decline of 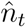 with the strength of interannual environmental fluctuations (*ϵ*, Fig. 5a) and in the fluctuations of *n*_*t*_ over time (i.e., 90% quantile of total population size, Fig. 5b). Sensitivity of 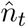 to fluctuations decreased among life histories as iteroparity 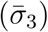 and maturation time 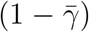 increased (see S1), from *pre-sem* to *pre-ite, del-sem*, and *del-ite* life histories. The standardization of the life histories for maximum growth rate 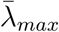 did not affect the rank order of life-history strategies. Overall, the sensitivity and the temporal fluctuations of *n*_*t*_ decreased when we increased 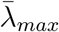 from 1.10 to 1.15 (Fig. 5a,b). A weaker strength of selection (*ω*^2^ = 16) also attenuated the sensitivity and stochasticity of population size for all life histories (Fig. S13).

#### Rate of directional trait evolution

As expected from our analytical predictions (Tab. S1), the biennial life cycle (*pre-sem*) had the fastest evolutionary trait response to directional environmental change, while the *del-ite* life history evolved the slowest (Fig. 5c). However, even though the differences were small, life histories standardized for 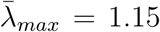 evolved faster than life histories with 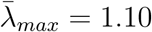 (Fig. 5c), which was a deviation from the analytical predictions. Nevertheless, differences in evolvability were mainly found on a yearly, absolute time scales. Once scaled to generation time, all life histories took a similar number of *generations* to reach the same phenotypic threshold (Fig. S12c).

Although standing genetic variation *G*_2_ was similar across life-history strategies after burn-in (Fig. 5d), its temporal dynamics was strongly affected by the degree of iteroparity and delayed maturation. *G*_2_ increased fastest during adaptation to new average local conditions in the *pre-sem* and slowest in the *del-ite* life cycle (Fig. 4c). Overall, larger interannual fluctuations tended to increase standing genetic variation in all life histories (Fig. 5d).

#### Extinction risk

The extinction risk of the population, evaluated as the total number of extinct replicates at a given time during the simulations (Fig. 6), varied among life histories. In general, population extinctions were delayed relative to the onset of directional environmental change and still occurred after the environmental change stopped (Fig. 6a,c). The less robust life-history strategies (e.g., *pre-sem*) experienced earlier extinction because of larger demographic stochasticity leading to earlier declines in *n*_*t*_ than robust life histories (e.g., *del-ite*; Fig. 4b,d). Nevertheless, the iteroparous life histories (*pre-ite, del-ite*) had larger extinction risks when we prolonged the period of environmental change from 200 to 800 years, at least at low rate of random environmental fluctuations (Fig. 6c). At high rates of interannual fluctuations, the bi-annual life cycle *pre-sem* always had the highest rate of extinction (Fig. 6b,d). Overall, extinction risks were attenuated by having larger fecundity when setting 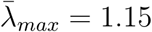 (Fig. S11) and when reducing the strength of selection to *ω*^2^ = 16, where no extinction occurred (Fig. S14).

**Figure 6:**
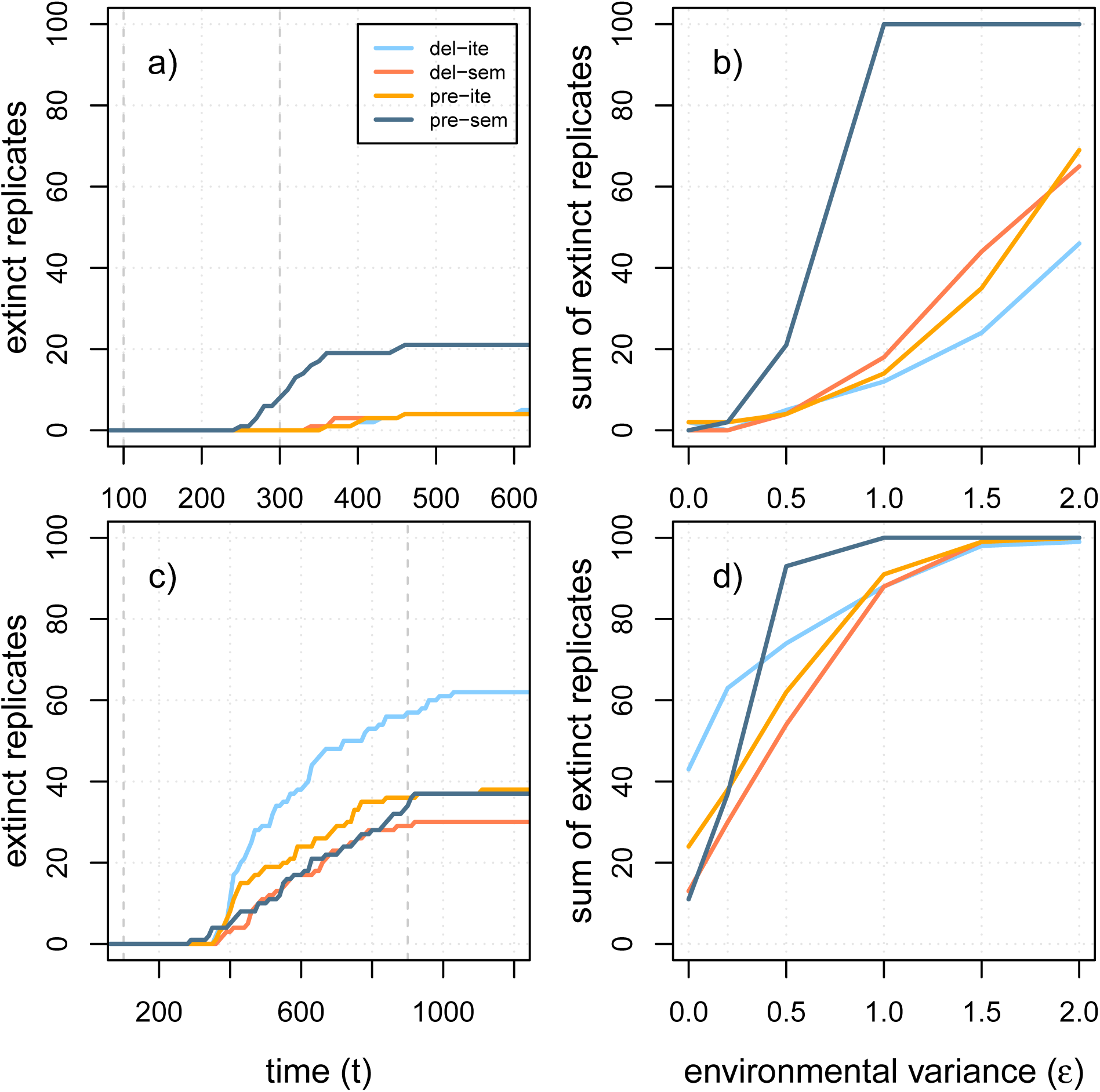
These graphs show the number of extinct replicates over time *t* (a,c) and the total number of extinct replicates depending on environmental variability *ϵ* (b,d) for each of the four life histories. Extinction dynamics are shown for two different scenarios with environmental change either lasting for 200 years (a,b) or for 800 years (c,d). The gray dashed lines indicate the start of environmental change (*t* = 100) and its end (*t* = 300 for a,b; *t* = 900 for c,d). In both scenarios, the rate of environmental change per year was set to 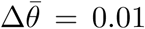 and all life histories were standardized for a maximum growth rate of 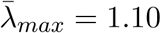. The temporal dynamics are shown for an environmental variability of *ϵ* = 0.5 (a) and *ϵ* = 0.2 (c).

## Discussion

Life-history strategies can impact a species’ persistence in a dynamic environment, especially by helping it to cope with short-term interannual fluctuations and by shaping its evolutionary response to long-term gradual changes (Lande, 1982; Hairston and De Stasio Jr., 1988; Barfield et al., 2011). In this study, we asked (i) whether a tradeoff between a life history’s demographic robustness against environmental variability and its rate of trait evolution exists, (ii) how life-history characteristics determine the position of a species along the robustness-evolvability tradeoff, (iii) whether a genetic storage effect could alleviate the tradeoff, and (iv) how the tradeoff affects species persistence in fluctuating environments. Using existing theory from matrix population modeling and quantitative genetics, we confirmed numerically that a tradeoff exists. However, current theory does not account for density-dependent population dynamics below carrying capacity and demographic feedback on the genetic variance of a population. For these reasons, we ran stochastic individual-based simulations. Here too, we could confirm the hypothesized tradeoff between a life history’s robustness and evolvability. Overall, we could show that, along the fast-slow continuum, short-lived semelparous species were more “evolvable” and could cope best with long-lasting directional environmental change, but experienced the highest extinction risk from interannual fluctuations and during the initial phase of environmental changes. They should thus suffer more from rapid anthropogenic climate changes than long-lived iteroparous species.

A higher robustness against random and uncorrelated between-year environmental fluctuations was achieved by three life-history characteristics: Offspring dormancy, higher adult survival (leading to iteroparity), and a larger maximum growth rate. Offspring dormancy and higher adult survival helped to tolerate a variable environment by shielding a higher proportion of individuals from detrimental environmental conditions and spreading the risk of failed survival across years (e.g., see also Orzack and Tuljapurkar, 1989; Tuljapurkar, 1990; Koons et al., 2008; Zeineddine and Jansen, 2009). In our simulations, offspring dormancy and iteroparity allowed the population to maintain larger sizes in face of fluctuations, while reducing its demographic stochasticity. In addition, a larger maximum growth rate improved the robustness to environmental variability by helping to recover faster from extreme events (Hallett et al., 2018).

Opposed to the means of promoting robustness, evolvability of life-history strategies benefited from lower dormancy in offspring and reduced adult survival, following two main effects. First, the rate of trait evolution in stage-structured populations generally increases when exposing more individuals to selection and improving their contribution to the next generation. From Equations 3 and 4, these two factors maximize the weight put on the amount of additive genetic variation *G*_2_ available for selection (i.e, the evolvability of the trait, see Houle, 1992; Hansen and Houle, 2008). Kuparinen et al. (2010) also showed this effect when simulating the evolution of a quantitative trait in Scots pine and Silver birch. A higher mortality in adult trees (e.g., from senescence or forest management) favored faster recruitment of adapted juvenile trees and accelerated trait evolution in their simulations. A second effect favoring evolvability was shorter generation times (also described in Barfield et al., 2011; Orive et al., 2017). Per unit of generation time, long-lived species experienced larger environmental shifts causing larger maladaptation and thus lower growth rate, eventually reducing juvenile recruitment rates sufficiently to cause larger extinction rates than for fast evolving, short-lived species. Osmond et al. (2017) described this mechanism in predator-prey systems when predation reduced the generation time of the prey species and thus helped the prey to better adapt to novel environmental conditions.

In our simulations, the genetic storage effect did not affect the amount of standing genetic variation at the start of directional environmental change and did not help to overcome the tradeoff between robustness and evolvability in long-lived species. Although all life histories had the same standing genetic variation, they differed markedly in the evolution of additive genetic variance *G*_2_ during environmental change. The genetic variance *G*_2_ increased during directional selection in all simulations, as expected from the increase of allele frequencies of rare variants during adaptation towards a new trait optimum (Bürger, 1999; Kopp and Matuszewski, 2014). This rise in *G*_2_ happened most rapidly for the biennial life cycle and slowest for the life history with offspring dormancy and iteroparity. Life histories thus shaped the rate of evolution via *G*_2_ differently from our initial expectations, in fact in the opposite direction, such that the tradeoff was not mitigated but enforced. Yet, the storage effect has been shown to speed up evolution with overlapping generations for very large levels of environmental fluctuations *ϵ* (Yamamichi et al., 2019). However, we could not impose such high levels of *ϵ* without driving populations to extinction when applying hard selection, while Yamamichi et al. (2019) simulated a yearly offspring production that was insensitive to *ϵ*.

To overcome the tradeoff, a higher maximum growth rate could allow life-history strategies to be more robust and at the same time more evolvable. In our models, a higher population growth rate 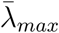 improved robustness against environmental fluctuations by helping to recover faster from detrimental environmental events and avoid extinction, in line with previous theory (Lynch and Lande, 1993; Lande, 1993; Bürger and Lynch, 1995; Bürger, 1999). Interestingly, a higher growth rate also caused faster trait evolution in our simulations, even though it had no or only very little effect on 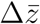 in our mathematical models. Here, a higher 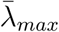, when realized by larger fecundity, was associated with more frequent mutation and recombination events. This effect of life-history parameters on the amount of mutational input (Fig. S10) is a process not accounted for in previous analytical models, but should be of importance in natural systems.

In conclusion, our results show that the feedback between evolutionary and demographic processes are critical for species persistence. A similar conclusion can be reached from models of evolutionary rescue (i.e., adaptation to single-shift environmental scenarios: Lynch et al., 1991; Lynch and Lande, 1993; Bürger and Lynch, 1995; Gomulkiewicz and Holt, 1995; Gomulkiewicz and Houle, 2009; Chevin, 2013), or from models of species persistence on temporally shifting environmental gradients (i.e., models of species’ range evolution: Polechová et al., 2009; Duputié et al., 2012). However, these models often ignore short-term perturbations (but see Orive et al., 2017), assume constant population growth (i.e., with density-independent growth, but see Boulding and Hay, 2001), and do not incorporate explicit life cycles. On the other hand, quantitative genetics models of trait evolution in stage-structured populations with (Engen et al., 2011, 2013) or without (Barfield et al., 2011) stochastic environmental fluctuations do not model species persistence, to a few recent exceptions (Marshall et al., 2016; Orive et al., 2017; Cotto et al., 2019). Importantly, no study to date explicitly addressed the tradeoff between evolvability and demographic robustness. We have shown that such a tradeoff exists and may limit the odds of evolutionary rescue in fast life histories, while slow life histories may persist longer while accumulating large adaptation lags at the cost of increased extinction debts (Dullinger et al., 2012; Cotto et al., 2017).

### Implications

We highlighted a fundamental property of eco-evolutionary dynamics whereby a larger evolutionary response in population adapting to a fluctuating environment comes at the expense of its demographic robustness to variation in vital rates. Our conclusions are based on comparisons of eco-evolutionary dynamics of standardized life cycles. However, life-history strategies in natural systems are not standardized. In fact, systematic differences in more than one life-history characteristic exist. Species with low generation times often exhibit systematically higher population sizes 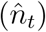, greater intrinsic growth rates, larger dispersal abilities, but lower competitive abilities than species with long generation times (Petit and Hampe, 2006; Buoro and Carlson, 2014; Beckman et al., 2018). They might thus be more robust to changes than predicted here. Similarly, long-lived species may suffer more from climate changes if adult mortality is increased, reducing their robustness but increasing their evolvability. For instance, adult trees may be sensitive to extreme weather events such as drought or forest fires (e.g., Van Mantgem and Stephenson, 2007). By simplifying the action of selection on a single life stage (e.g., on tree seedlings), we have not incorporated possible feedback between environmentally induced selection and life history characteristics. Such feedback could change the life history of the species, as shown by Cotto et al. (2019), and move the species along the fast-slow life history tradeoff between evolvability and robustness. Instead, we have characterized archetype life-history strategies that span the fast-slow continuum and exemplified those possible outcomes. Our work thus shows the importance of merging population ecology with evolutionary quantitative genetics in a dynamical and stochastic modelling framework to characterize species’ vulnerability to environmental changes.

## Acknowledgements

We thank Thomas F. Hansen, Luis-Miguel Chevin, Olivier Cotto, and Richard Gomulkiewicz for insightful discussions and comments on the manuscript. MS and FG were supported by grants PP00P3 144846 and PP00P3 176965 from the Swiss National Science Foundation to FG.

## Appendix

### Appendix A: 3-stage model

#### Equilibrium population size

The populations’ demographic dynamics were modeled via the population vector

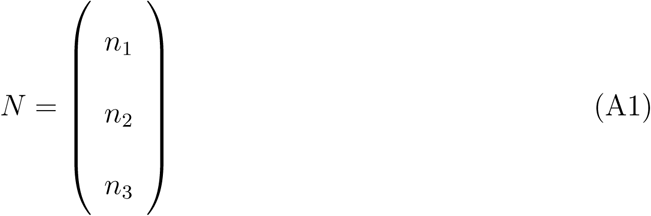

and the matrix population model

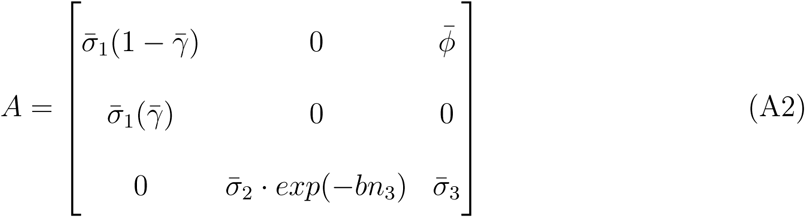

such that *N*(*t* + 1) = *AN*(*t*). By solving for the stage densities at equilibrium when 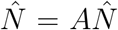,with 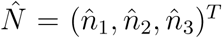, the individual numbers within the offspring, juvenile, and adult stage at equilibrium could be derived as

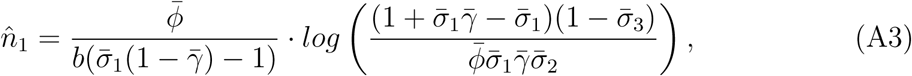

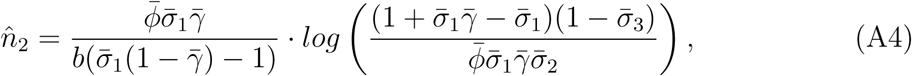

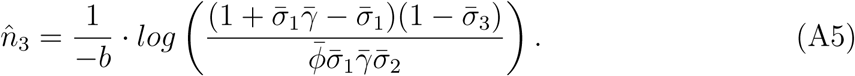

The total population size at equilibrium was computed from 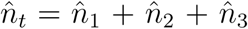 as

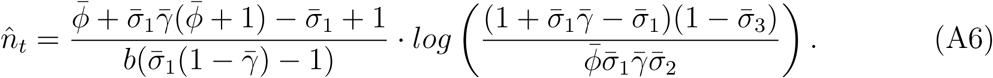

#### Sensitivity of total population size to juvenile survival

The partial derivative of the total population size at equilibrium 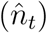 to juvenile survival 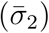 was calculated as

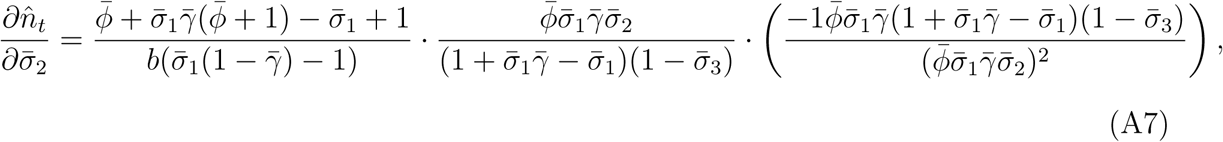

which could be simplified to

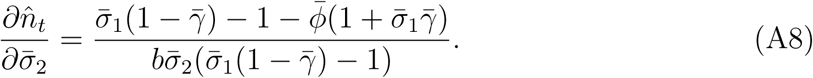

To standardize a life history for a specific total equilibrium population size 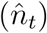 the strength of intra-specific competition was adjusted as follows

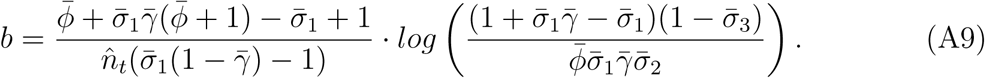

By inserting equation A9 in equation A8, the population size sensitivity can be expressed depending on the equilibrium population size as

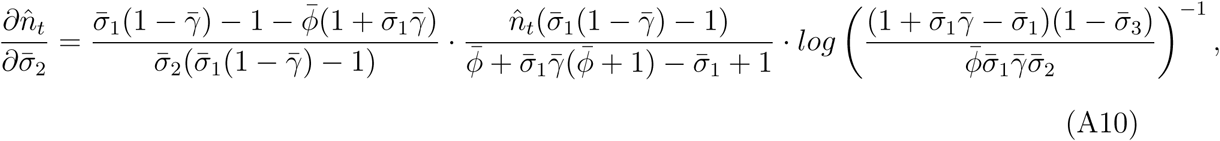

such that the following partial derivative is obtained

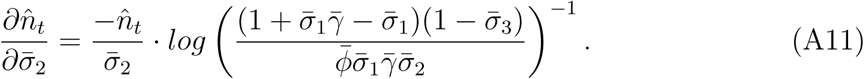

#### Rate of evolution with exponential growth

We adapted equation 8 from Barfield et al. (2011) for our three-stage life cycle in absence of density regulation as follows. We assume that the vital rate of juveniles can be expressed as 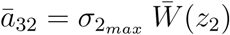, the product of maximum juvenile survival 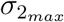 and average juvenile survival depending on the match between phenotype and environment 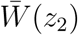. Then, with *ā*_32_ 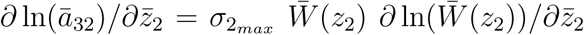, and writing 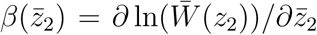 (Lande, 1979; Lande and Arnold, 1983) as well as 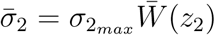, the asymptotic rate of evolution is

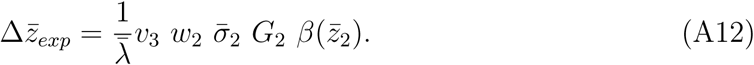

The juvenile proportion (*w*_2_) could be derived from the right eigenvector of *A* (Caswell, 2001, p. 87) when solving 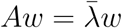 as

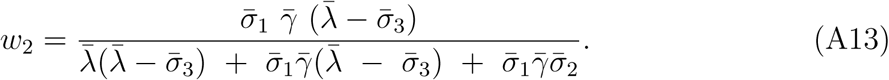

The reproductive value of adults (*v*_3_) could be derived from the left eigenvector (*v*) of *A* when solving 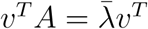 (Caswell, 2001, equation 4.81) as

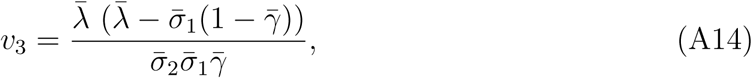

and normalized to

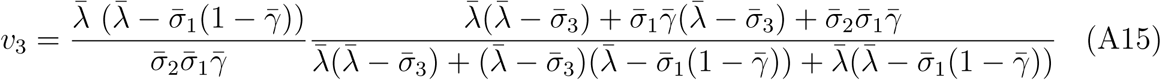

such that *v* ∗ *w* = 1 (as in Barfield et al., 2011). The rate of evolution depending on 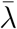 becomes:

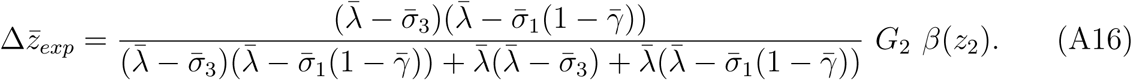

#### Rate of evolution at equilibrium population size

With density-dependent population growth, we assume that the rate of evolution can be approximated from the Lande theorem as well (Barfield et al., 2011), at least in absence of density-dependent selection when the population reached equilibrium density 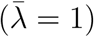, and when population dynamics are faster than evolutionary dynamics. Then, competition survival *c* and the transition matrix *A* are nearly constant over time and the population reached SSD, while not being at the phenotypic optimum yet. Furthermore, the growth rate sensitivity 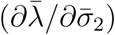 of both models, with exponential growth and at equilibrium population size, is then equivalent (see Figure S9).

Equation 8 from Barfield et al. (2011) describes the rate of evolution 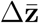 depending on the asymptotic growth rate 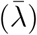, single vital rates (*ā*_*ij*_), stage proportions (*w*_*j*_), reproductive values (*v*_*i*_), additive genetic variances (*G*_*j*_), and the gradient operator 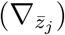 as

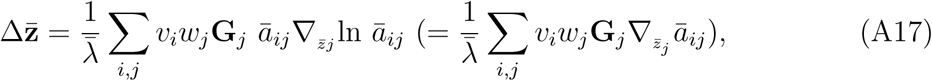

which we adapted for our three-stage life cycle with only juveniles under selection. Here, we assume that the vital rate of juveniles can be expressed as 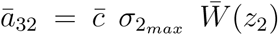, the product of maximum juvenile survival 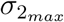, average juvenile survival depending on the match between phenotype and environment 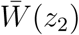, and competition survival 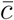. Then, with 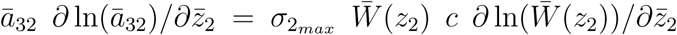, and writing 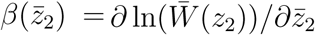 (Lande, 1979; Lande and Arnold, 1983) as well as 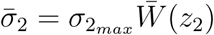, the asymptotic rate of evolution is equal to

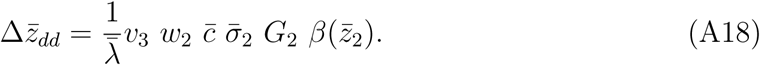

The juvenile proportion (*w*_2_) could be derived from equation A4 and A6 as

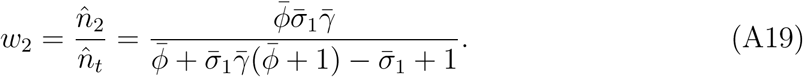

The probability of surviving intra-specific competition 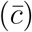 was derived by combining the Ricker function 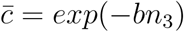 and equation A5 as

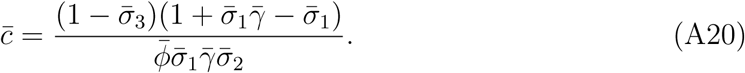

The reproductive value of adults (*v*_3_) could be derived from the left eigenvector (*v*) of the matrix population model (*A*) when solving 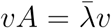 (Caswell, 2001, equation 4.81). For a population at equilibrium population size 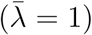 the reproductive value of adults can be derived as

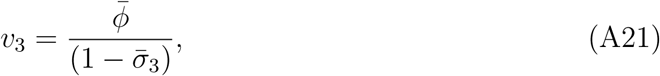

and normalized to

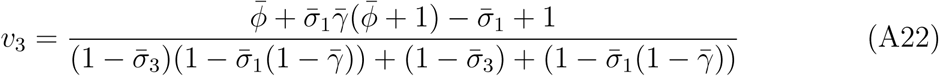

such that *v* ∗ *w* = 1 (as in Barfield et al., 2011). The rate of evolution at equilibrium population density 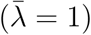 becomes:

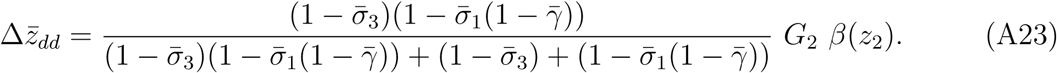

### Appendix B: 2-stage model

#### Equilibrium population size

The demographic dynamics for a 2-stage life history with non-overlapping generations were modeled by the population vector

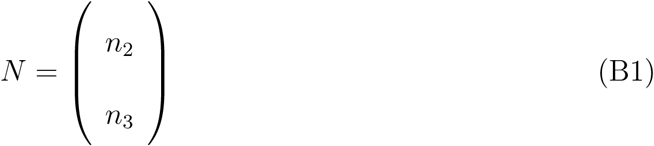

and the matrix population model

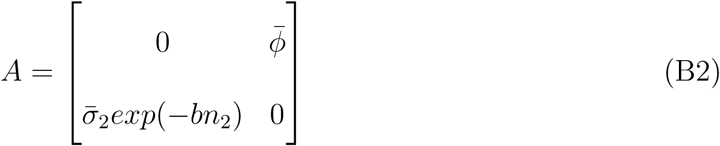

such that *N* (*t* + 1) = *AN*. By solving for the individual numbers within stages at equilibrium when 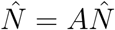 and 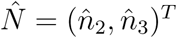 the juvenile and adult numbers at equilibrium were derived as

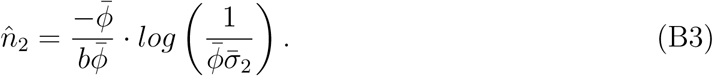

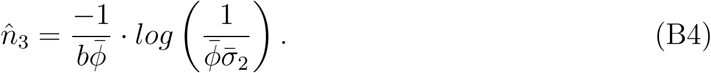

(with natural logarithm log()). The total population size at equilibrium was computed from 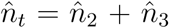 as

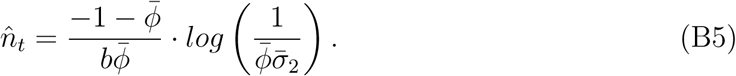

#### Sensitivity of total population size to juvenile survival

The partial derivative of the total population size at equilibrium 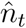 to juvenile survival 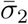 was derived as

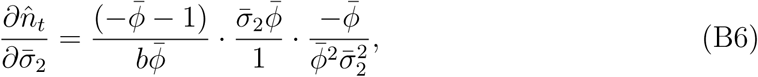

which could be simplified to

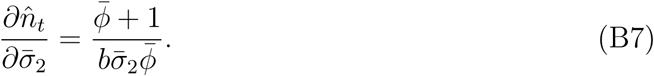

To reach a certain total equilibrium population size 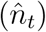 the strength of intra-specific competition (*b*) was standardized as follows

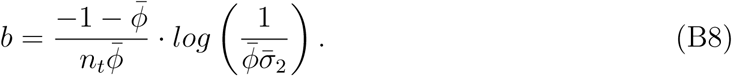

Given that the maximum growth rate 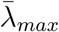 could be calculated from

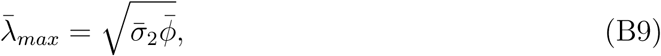

life histories could be standardized for 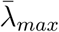 via 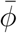 adjustments as

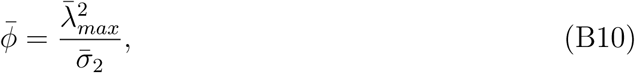

such that the sensitivity 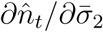 could be generalized to

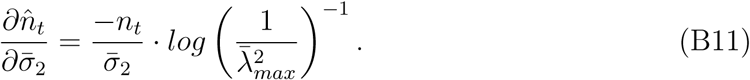

#### Rate of evolution with exponential growth

To describe the rate of evolution we adapted equation 8 from Barfield et al. (2011) for our two-stage life cycle similar to the three-stage life history as

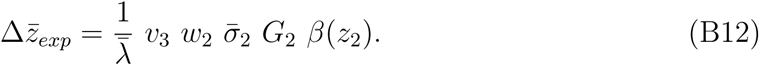

The proportion of juveniles (*w*_2_) could be derived from the right eigenvector of *A* as

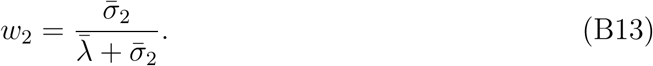

The reproductive value of adults (*v*_3_) could be derived from the left eigenvector (*v*) of *A* as

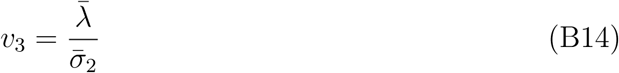

and normalized to

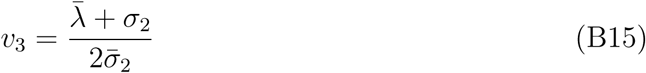

such that *v* ∗ *w* = 1 (as in Barfield et al., 2011). The rate of evolution depending on the single vital rates then could be expressed as

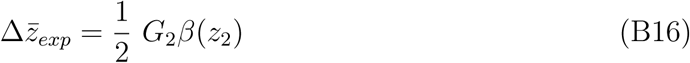

#### Rate of evolution at equilibrium population size

To describe the rate of evolution at carrying capacity 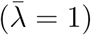 we adapted equation 8 from Barfield et al. (2011) for our two-stage life cycle similar to the three-stage life history as

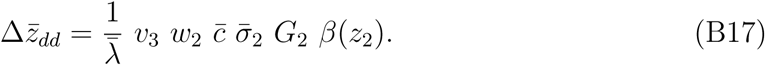

The juvenile proportion could be derived from equation B3 and B5:

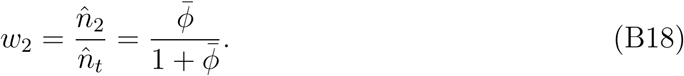

The probability of surviving intra-specific competition (*c*) was derived by combining the Ricker function 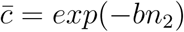 and equation B3:

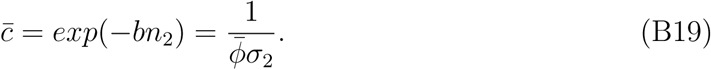

The reproductive value of adults (*v*_3_) could be derived from the left eigenvector (*v*) of the matrix population model (*A*) when solving 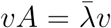 (Caswell, 2001, equation 4.81). For a population at equilibrium 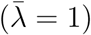 the reproductive value of adults could be derived as

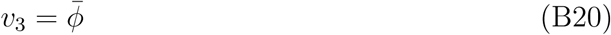

and normalized to

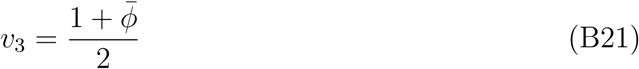

such that *v* ∗ *w* = 1 (as in Barfield et al., 2011). The rate of evolution depending on the single vital rates then could be expressed as

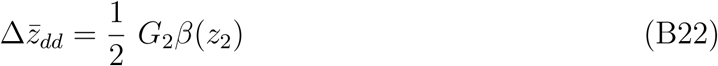

## Supplementary Material

## Supp A: 3-stage model with Beverton-Holt density regulation

### Equilibrium population size

The demographic dynamics were modeled via the population vector

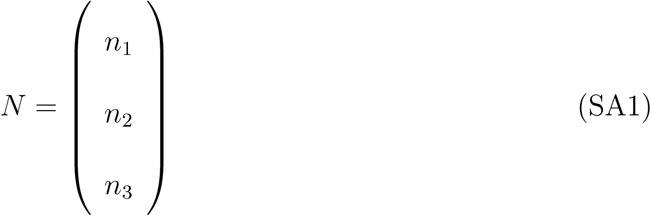

and the MPM

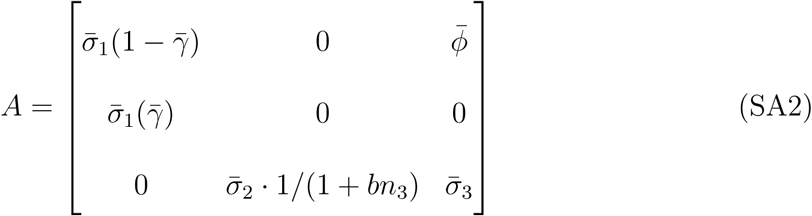

such that *N* (*t* + 1) = *AN* (*t*). At equilibrium 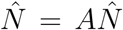 with 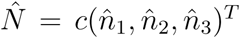. The number of offspring, juveniles, and adults at equilibrium then could be derived as

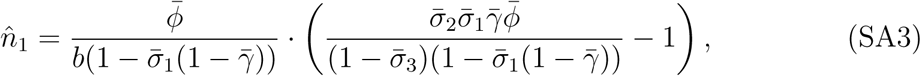

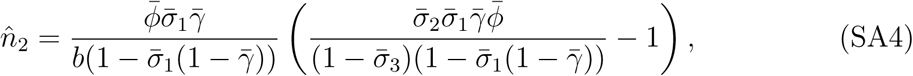

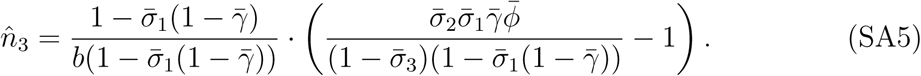

The total population size at equilibrium was computed from 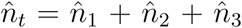 as

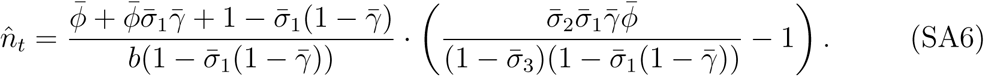

### Sensitivity of total population size to juvenile survival

The partial derivative of the total population size at equilibrium 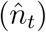 to juvenile survival 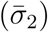 was calculated as

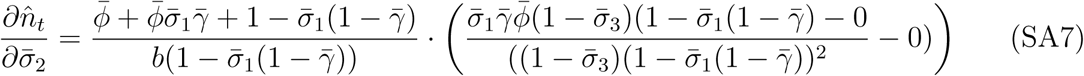

and could be simplified to

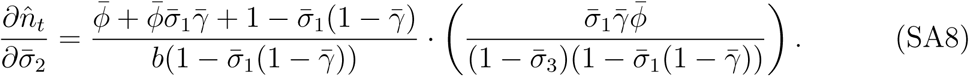

To reach a certain total equilibrium population size 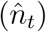 the strenght of intra-specific competition can be standardized as follows

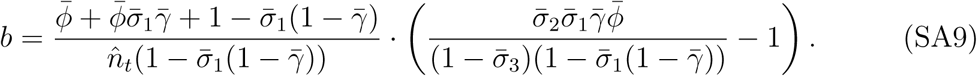

By inserting equation SA9 in SA8 the population size sensitivity to juvenile survival could be standardized for 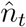 as

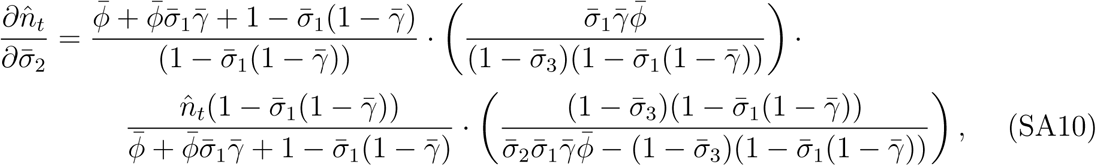

such that the following partial derivative is obtained

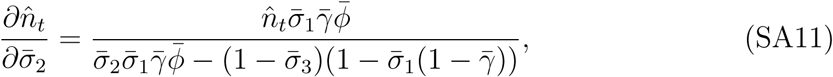

### Rate of evolution with exponential growth

The asymptotic rate of trait evolution in absence of density regulation is independent of the competition model as 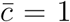 such that 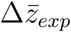 for Beverton-Holt density regulation is identical to 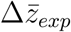 with the Ricker model in Appendix A (equation A16).

### Rate of evolution at stable population size

The equation from Barfield et al. (2011) could be adapted for our 3-stage life cycle at equilibrium population density 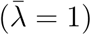 as

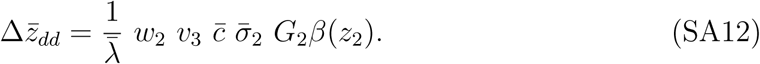

The juvenile proportion (*w*_2_) can be derived from equation SA4 and A6 as

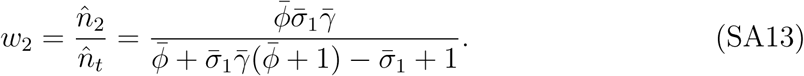

The probability of surviving intra-specific competition 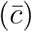 can be derived by combining the Beverton-Holt function 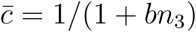 and equation SA5

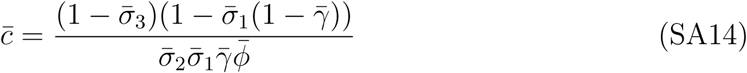

The reproductive value of adults (*v*_3_) can be derived from the left eigenvector (*v*) of the matrix population model (*A*) when solving 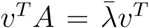 (Caswell 2001, chapter 4.6, equation 4.81). For a population at its stable population size 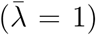 the reproductive value of adults is

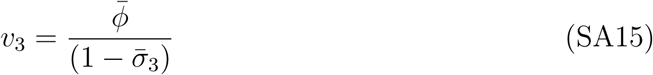

The normalization of *v*_3_ such that *v*^*T*^ *w* = 1 could be achieved as

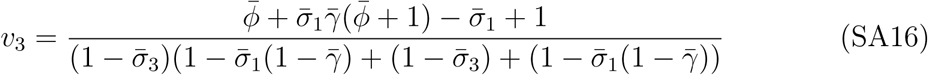

The rate of evolution then can be expressed as

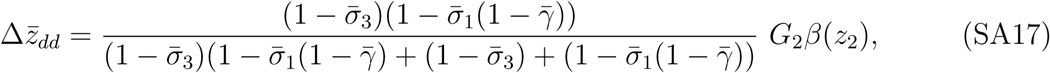

which is identical to 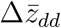 for the Ricker model (equation A23).

## Supp B: 2-stage model with Beverton-Holt density regulation

### Equilibrium population size

Population dynamics were modeled with the population vector

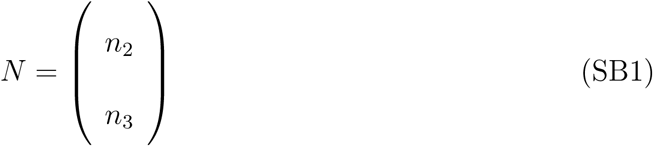

and the matrix population model

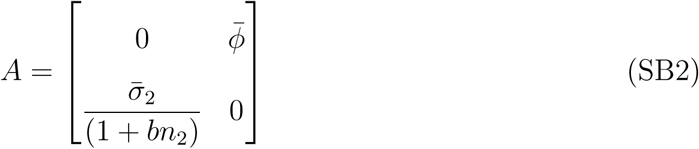

such that *N* (*t* + 1) = *AN* (*t*) with equilibrium 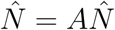.

The number of juveniles and adults at equilibrium then could be derived as

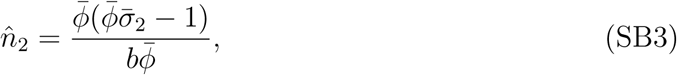

and

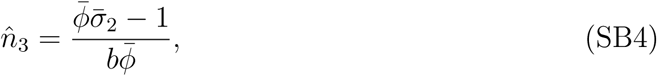

with a total population size at equilibrium of

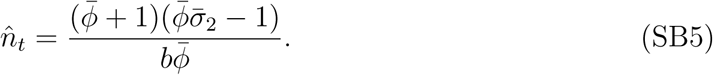

### Sensitivity of total population size to juvenile survival

The partial derivative of the total population size at equilibrium 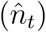 with respect to juvenile survival 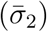 can be computed as

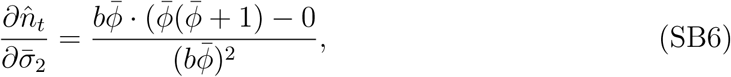

and simplified to

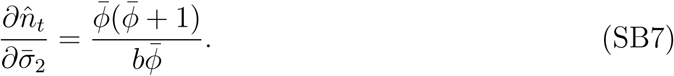

The strength of intra-specific competition (*b*) could be standardized to reach a certain equilibrium population size 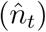 via

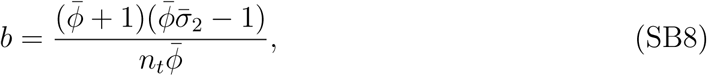

while the fecundity 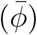 could be used to standardize a life history for a specific maximum growth rate 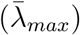 when

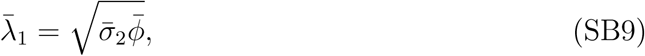

and

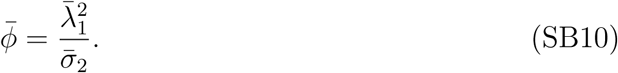

The sensitivity 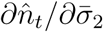 then could be reformulated as

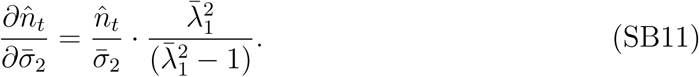

### Rate of evolution with exponential growth

The asymptotic rate of trait evolution in absence of density regulation is independent of the competition model (Ricker or Beverton-Holt) as 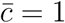 such that 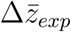 for the 2-stage model is identical to Appendix B (equation B16).

### Rate of evolution at stable population size

The equation from Barfield et al. (2011) can be adapted for our two-stage life cycle as

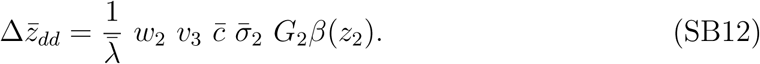

The juvenile proportion can be derived from equation B3 and B5:

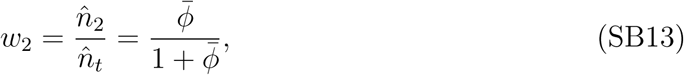

and the probability of surviving intra-specific competition (*c*) can be derived by combining the Beverton-Holt function 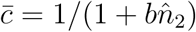 and equation B3:

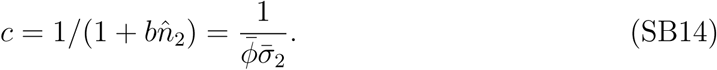

The reproductive value of adults for the 2-stage life history with Beverton-Holt density regulation is

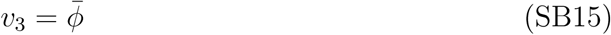

and can be normalize (*v*^*T*^ *w* = 1) to

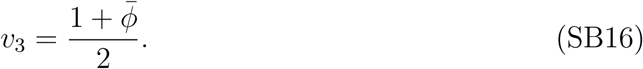

The rate of evolution for two-stage life history at population size equilibrium with BH density regulation and 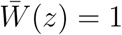 then can be expressed as

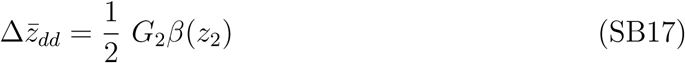

which is identical to 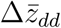 with the Ricker competition model (equation B22).

## Supp C: Plots mathematical model

**Figure S1:**
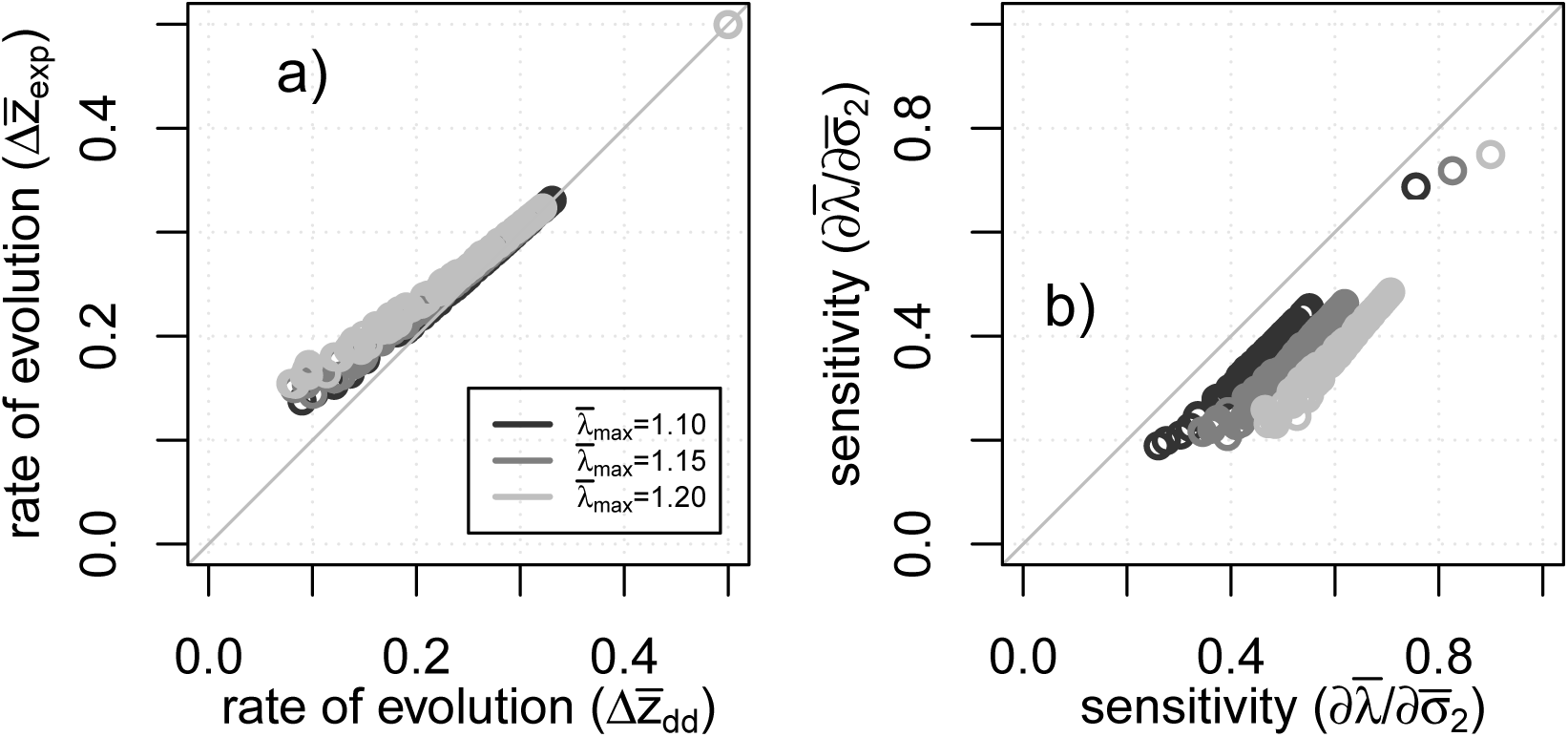
comparisons between exp. growth and density-dependent growth. The model results for the the rate of evolution 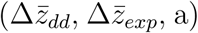 and the growth rate sensitivity 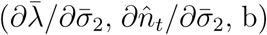 are plotted for both models with exponential growth (y-axis) and with density regulation (x-axis). Each circle illustrates a random combination of vital rates (with 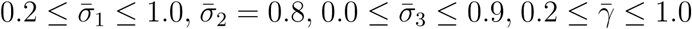). Fecundity 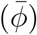 was re-calculated for each vital rate combination to standardize the maximum population growth rate 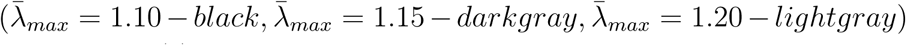. The strength of competition (*b*) was re-calculated for each life-history variant to standardize for the same total population size (*n*_*t*_=1’500) respectively.

**Figure S2:**
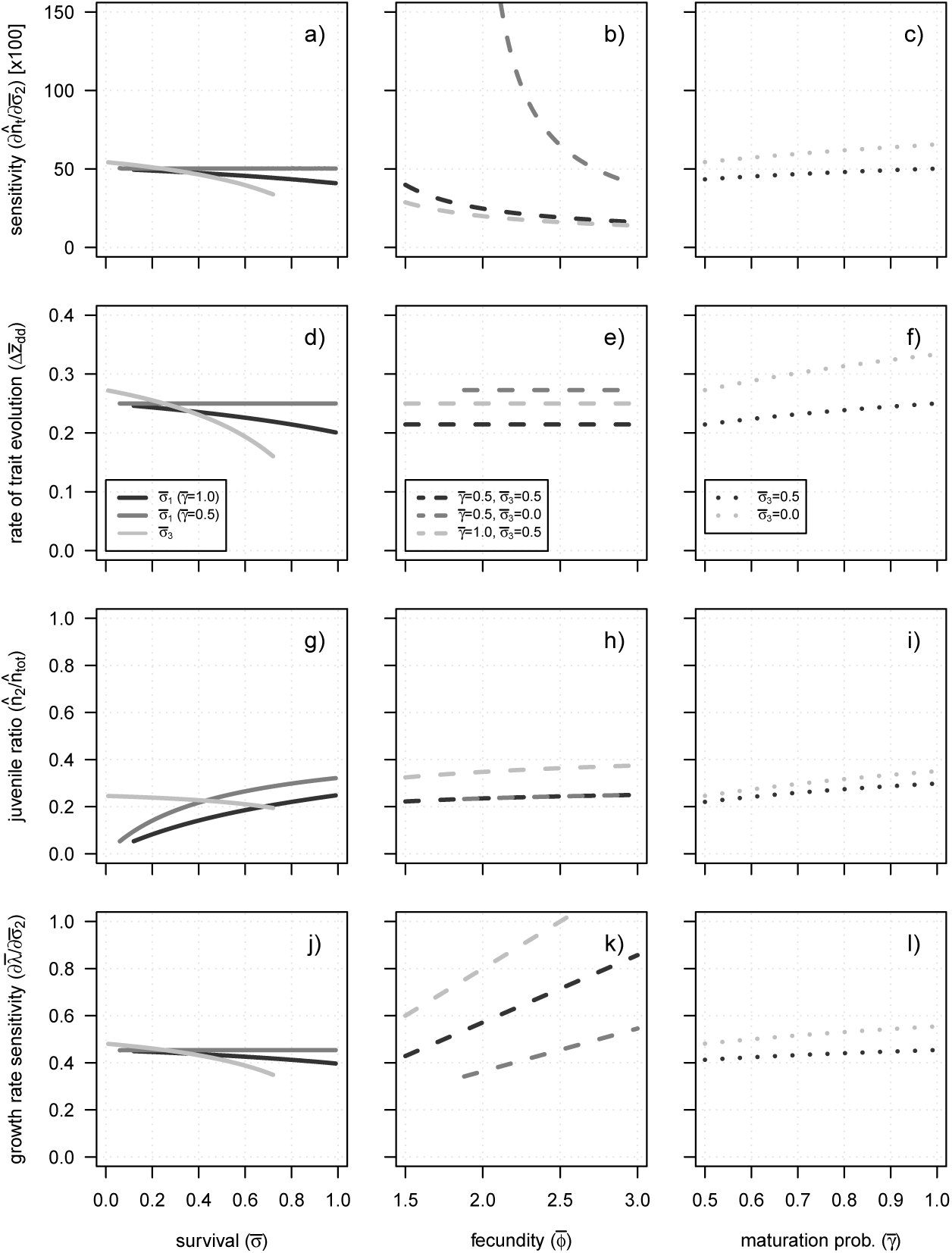
vital rate effects on robustness and evolvability (3-stage model) The model results for the sensitivity of the total population size (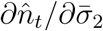, a-c), the rate of evolution (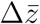, d-f), the juvenile ratio (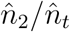, g-i), and the sensitivity of the max. growth rate (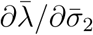, j-l) are shown depending on a life histories’ survival rate 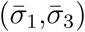, fecundity 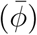, and maturation probability 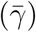. The maximum growth rate has been standardized for 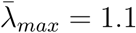 by fecundity adjustments, except for the case when we varied fecundity (b,e,h,k). Each parameter combination has been standardized for the same equilibrium population size in absence of interannual environmental fluctuations 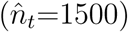 via adjustments of the strength of competition (**b**). The following parameter values have been used if not other specified: 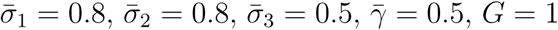, and *β*(*z*_2_) = 1. This graph is identical to figure 3 but shows additionally the juvenile ratio.

**Figure S3:**
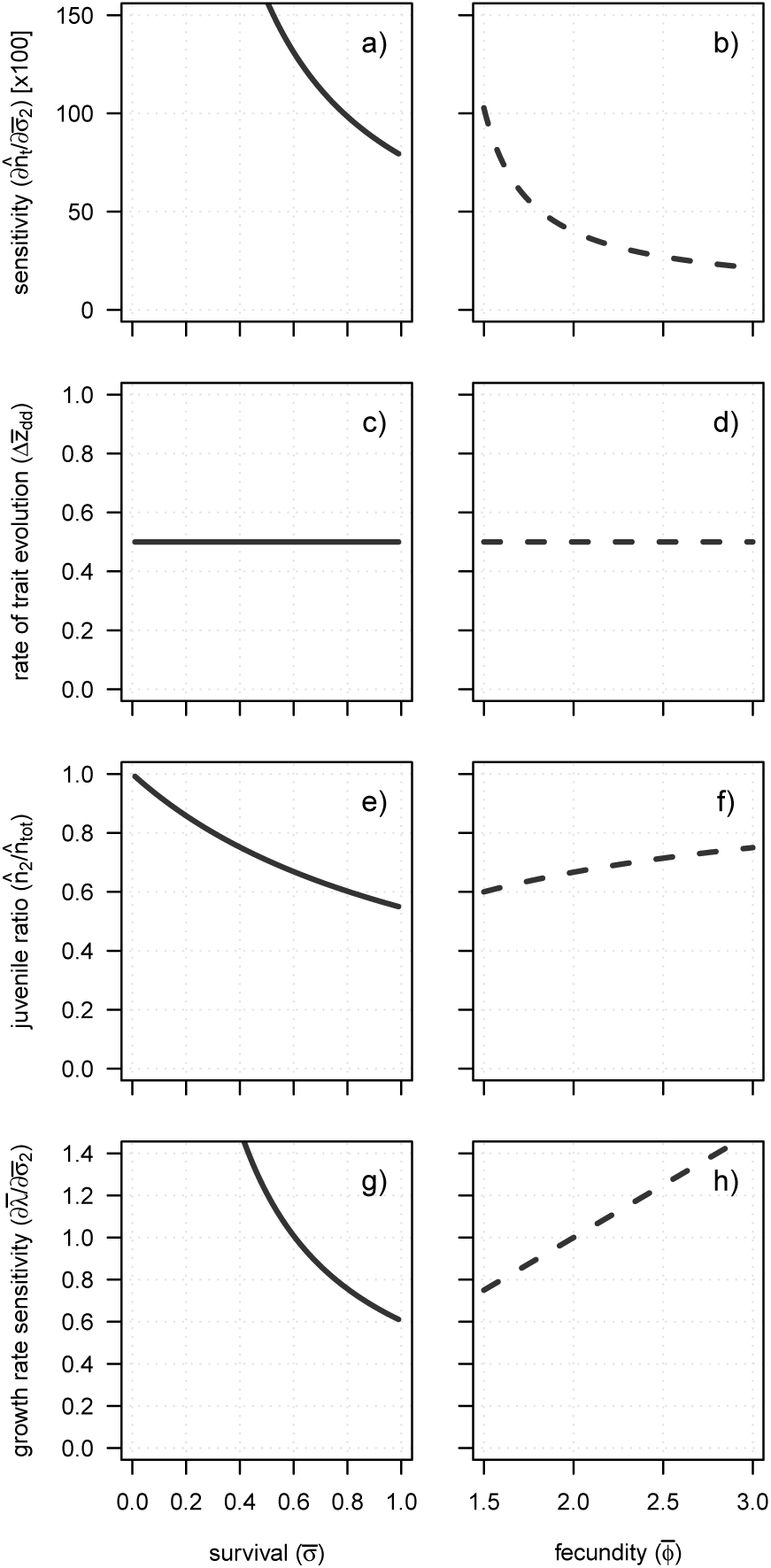
vital rate effects on robustness and evolvability (2-stage model) The sensitivity of the total population size (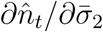, a-b), the rate of evolution (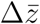, c-d), the juvenile ratio (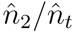, e-f), and the sensitivity of the max. growth rate (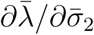, g-h) are shown depending on the two-stage life cycles’ survival rate 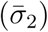 and fecundity 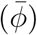. The maximum growth rate has been standardized for 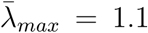 by fecundity adjustments, except for the case when we varied fecundity (b,d,f,h). Each parameter combination has been standardized for the same equilibrium population size in absence of interannual environmental fluctuations 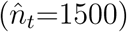 via adjustments of the strength of competition (**b**)., *G* = 1, and *β*(*z*_2_) = 1

**Figure S4:**
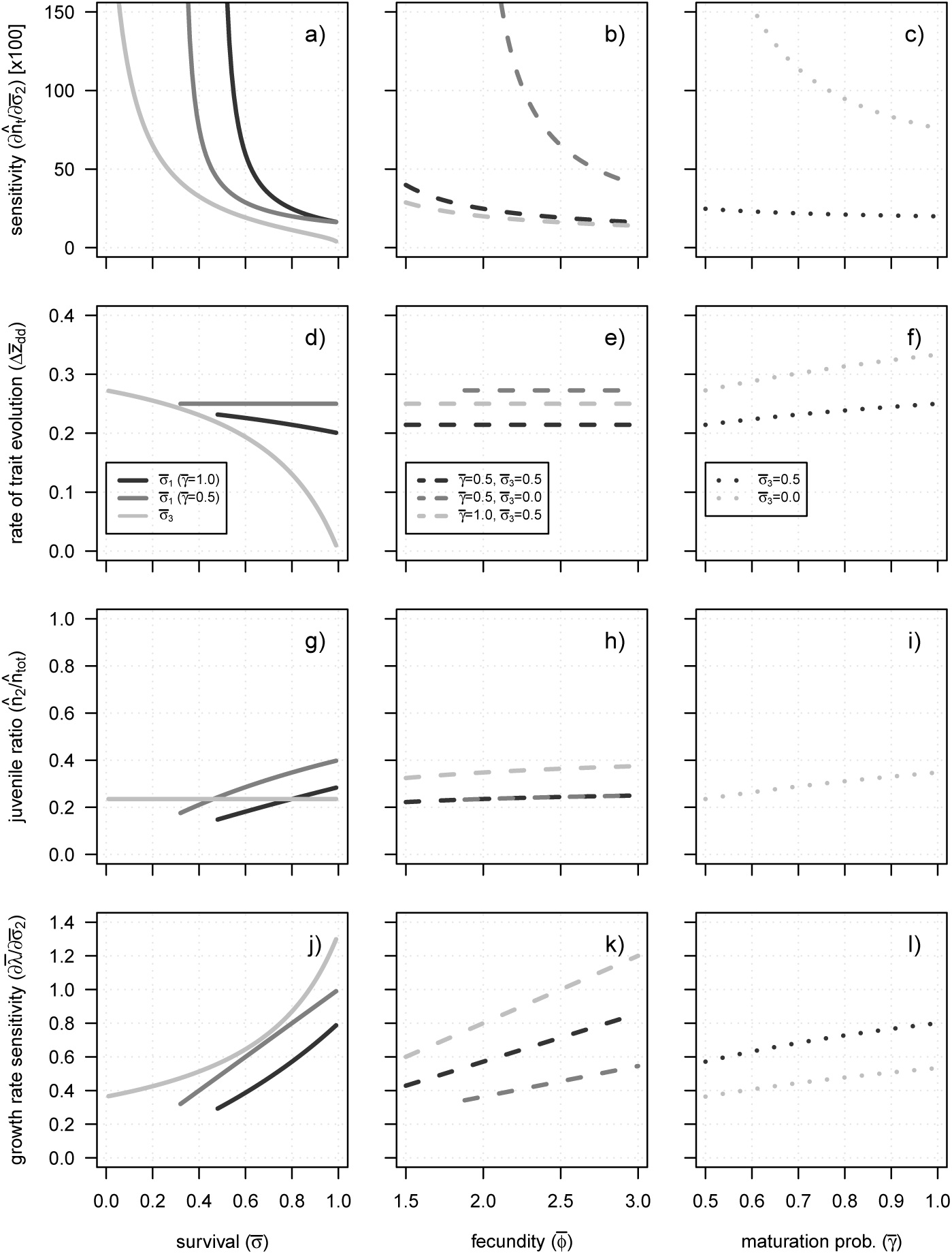
in absence of growth rate standardization (3-stage model) The model results for three-stage life cycles are shown for the sensitivity of the total population size (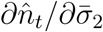, a-c), the rate of evolution (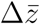, d-f), the juvenile ratio (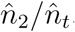, g-i), and the sensitivity of the max. growth rate (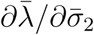, j-l) depending on a life histories’ survival rate 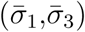, fecundity 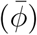, and maturation probability 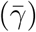. Each parameter combination has been standardized for the same equilibrium population size in absence of interannual environmental fluctuations 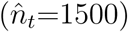 via adjustments of the strength of competition (**b**). However, the maximum growth rate has NOT been standardized in this case. The following parameter values have been used if not other specified: 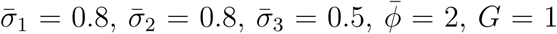, and *β*(*z*_2_) = 1. This figure shows the same results as figure 3 but adds the proportion of juveniles (g-i).

**Figure S5:**
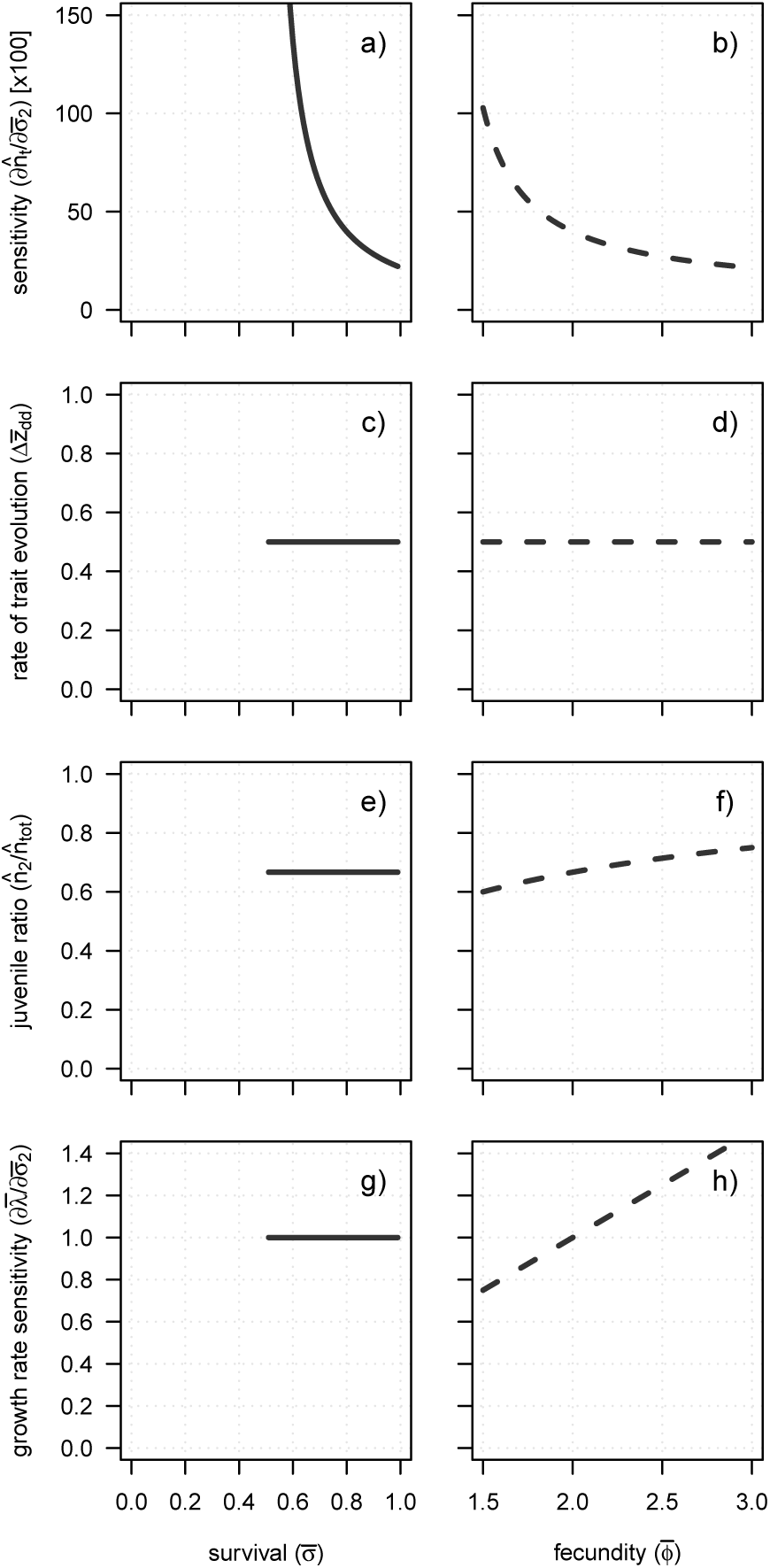
in absence of growth rate standardization (2-stage model) The model results for two-stage life cycles illustrate the sensitivity of the total population size (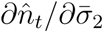, a-b), the rate of evolution (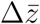, c-d), the juvenile ratio (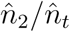, e-f), and the sensitivity of the max. growth rate (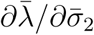, g-h) depending on a life histories’ survival rate 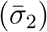 and fecundity 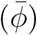. Each parameter combination has been standardized for the same equilibrium population size in absence of interannual environmental fluctuations 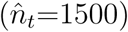 via adjustments of the strength of competition (**b**). However, the maximum growth rate has NOT been standardized for 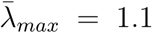 by fecundity adjustments. Juvenile survival was fixed to 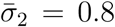 with varying fecundity (b,d,f,h) and fecundity was set to 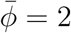 with varying 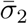 (a,c,e,g).

**Figure S6:**
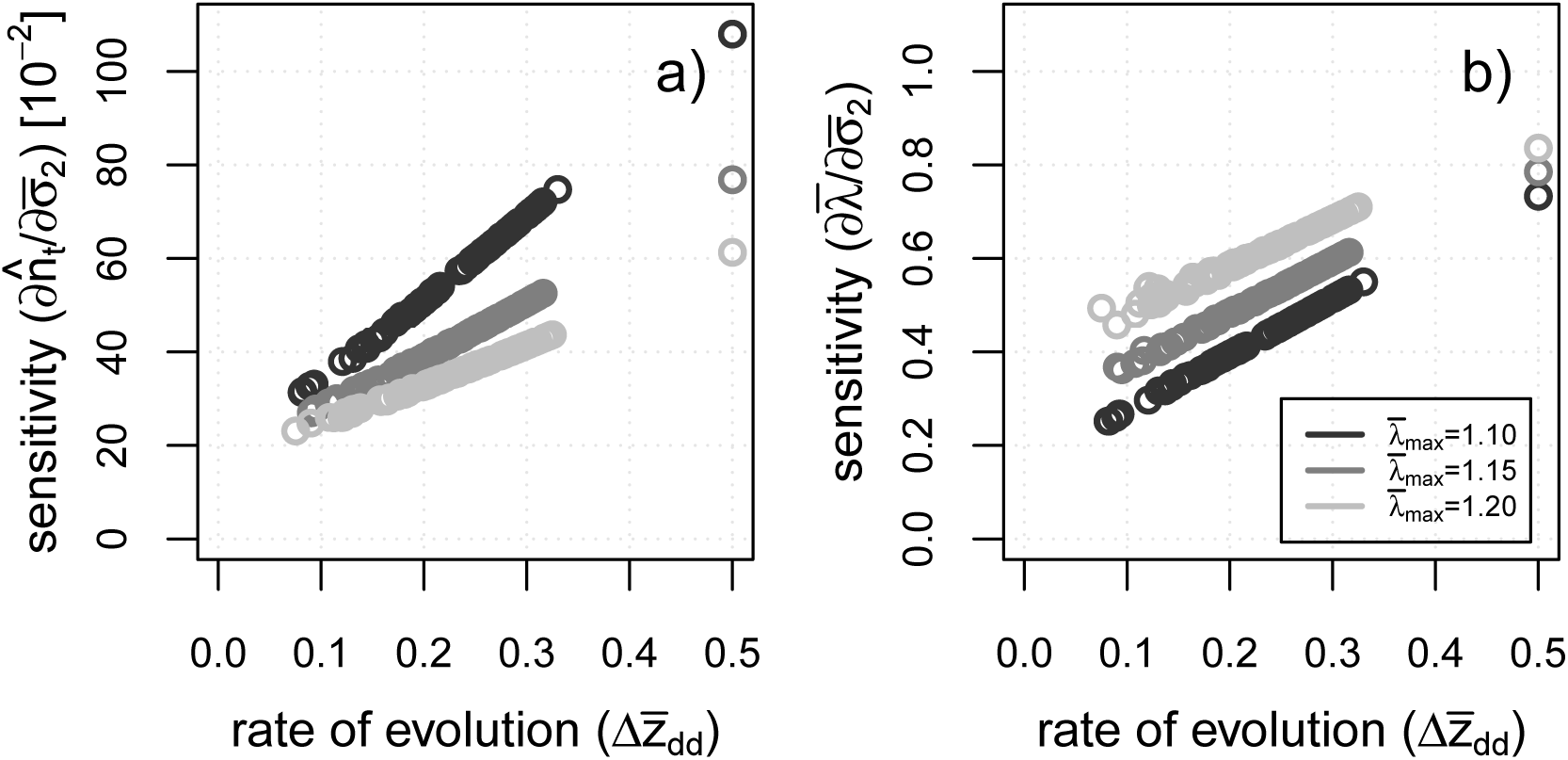
evolvability-robustness tradeoff with the Beverton-Holt model. A life history’s tradeoff between evolvability 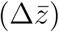 and robustness (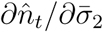 in a, and 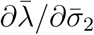 in b) using a Beverton-Holt function for density regulation. Each circle illustrates a random combination of vital rates (with 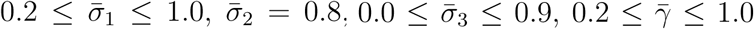). The fecundity 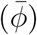 and the strength of competition (*b*) were re-calculated for each vital rate combination to standardize the maximum population growth rate 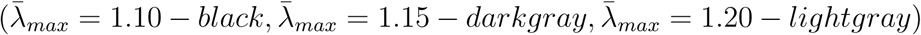 and the total population size at stable conditions (*n*_*t*_=1’500). We then calculated the rate of evolution (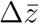, with *G*_2_*β*(*z*_2_) = 1), the population size sensitivity 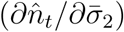, and the growth rate sensitivity 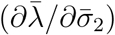 for each life-history variant. The three single points on the right of the plots were derived for the 2-stage life cycle (with 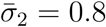), while the rest of the points illustrate parameter combinations for 3-stage life cycles.

**Figure S7:**
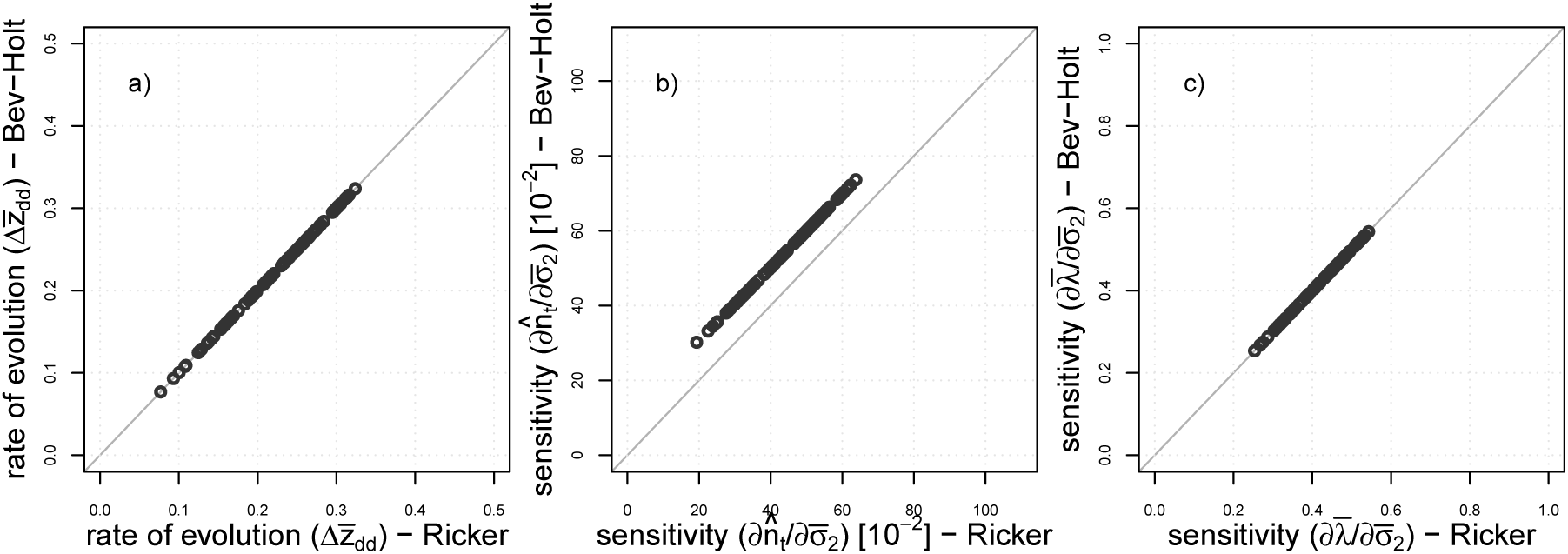
direct comparisons between Beverton-Holt and Ricker model. These graphs compare the model results for density regulation based on the Ricker function with the models based on the Beverton-Holt function, contrasting the rate of evolution 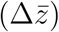, the sensitivity of the total population size to juvenile survival 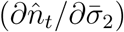, and the sensitivity of the population growth rate to juvenile survival 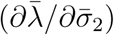. Each dot is a random combination of vital rates with 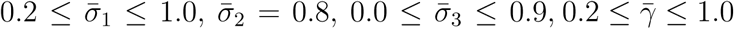.

**Figure S8:**
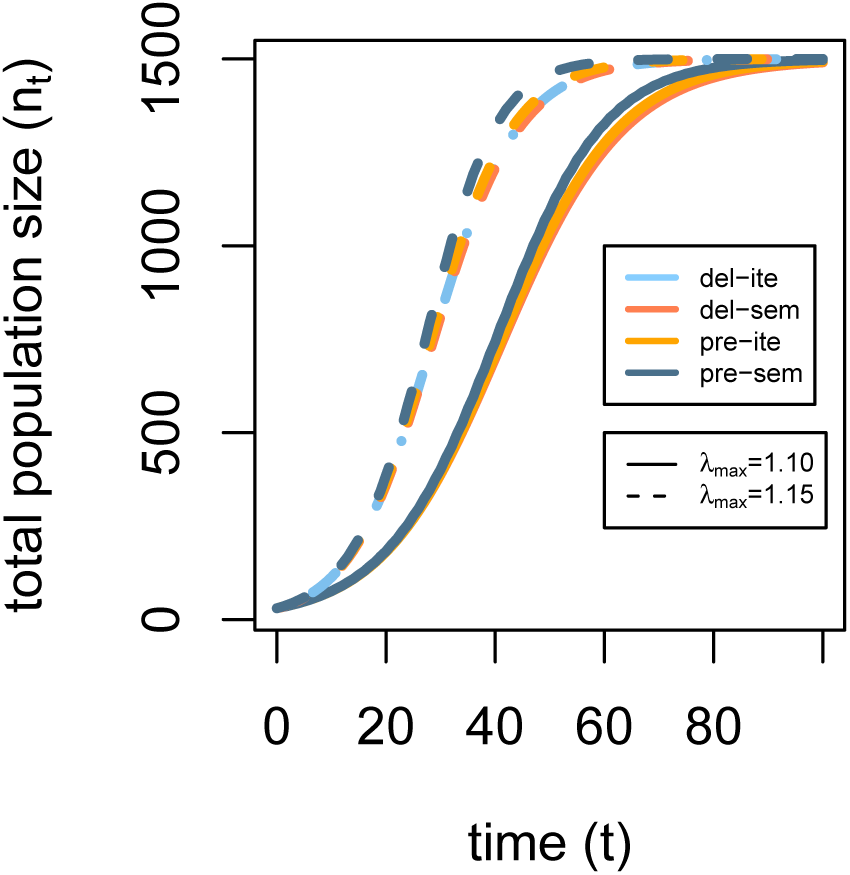
population growth for the simulated life histories. This graph illustrates the growth trajectories for all four life histories (*delite, del-sem, pre-sem, pre-ite*) and both maximum growth rates (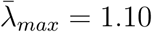 and 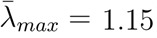) over time. All life histories here start at the same total population size (*n*_*t*_ = 30) and grow to the same carrying capacity of 1500 individuals (*n*_*t*_ = *n*_1_ + *n*_2_ + *n*_3_). The population growth trajectories are quite similar across life histories with the same 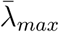, while 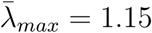 results in a faster increase in *n*_*t*_ than a standardization for 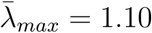. In the beginning, stage structure was adjusted to the SSD of the respective life history.

**Figure S9:**
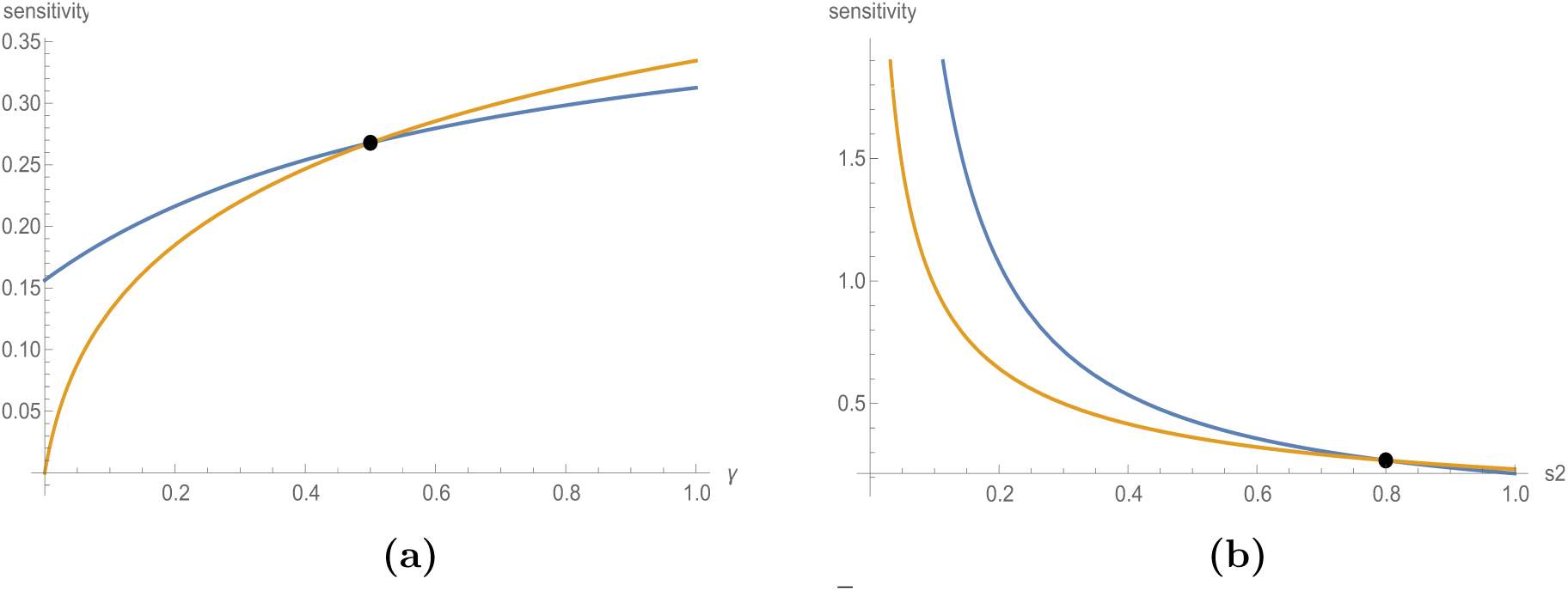
convergence of growth rate sensitivities. The growth rate sensitivity 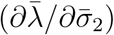 is compared between exponentially growing populations (yellow lines) and populations at carrying capacity (blue lines) depending on maturation probability 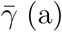, and juvenile survival 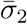 (b). The two black dots illustrate correspondence between both models at 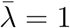. If not specified differently, vital rates were set to 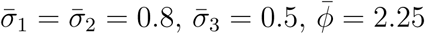, and 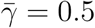.

## Supp D: Plots and table of the individual-based simulations

**Table S1:**
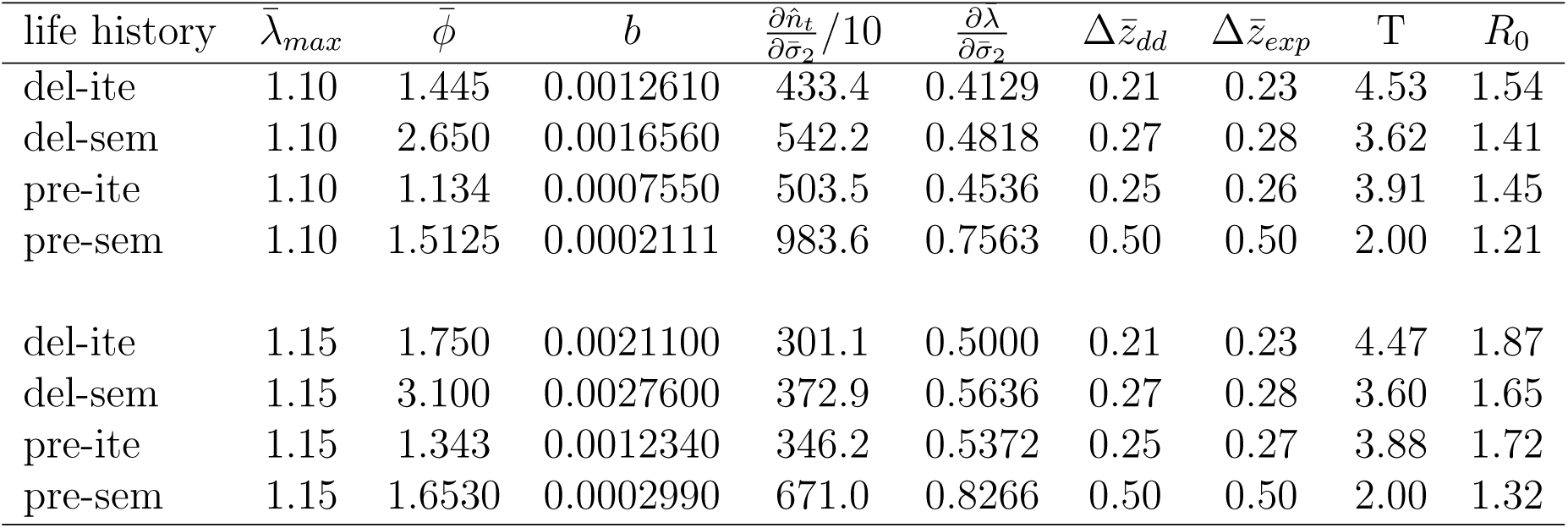
parameters of simulated life histories. The following table lists the vital rates of the simulated life-history strategies, which were combinations of delayed (*del*, maturation probability 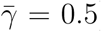) or precocious (*pre*, 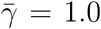) offspring maturation, and semelparous (*sem*, adult survival 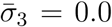) or iteroparous (*ite*, 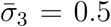) adults, standardized for 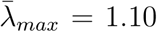 or 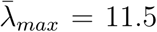 via the annual fecundity 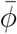. Offspring and juvenile survival were set to 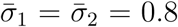 for all life histories. The generation time (*T*) and the net reproductive rate (*R*_0_) were calculated for the MPMs in absence of density regulation using the functions generation.time() and net.reproductive.rate() from the popbio package in R (Stubben and Milligan, 2007; R Core Team, 2019). The growth rate sensitivity 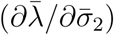 was calculated for the MPMs at 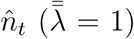 using explicit derivations from Mathematica 12.1 (Wolfram Research, 2020). The rate of evolution 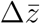 was calculated with *G*_2_*β*(*z*_2_) = 1.

**Figure S10:**
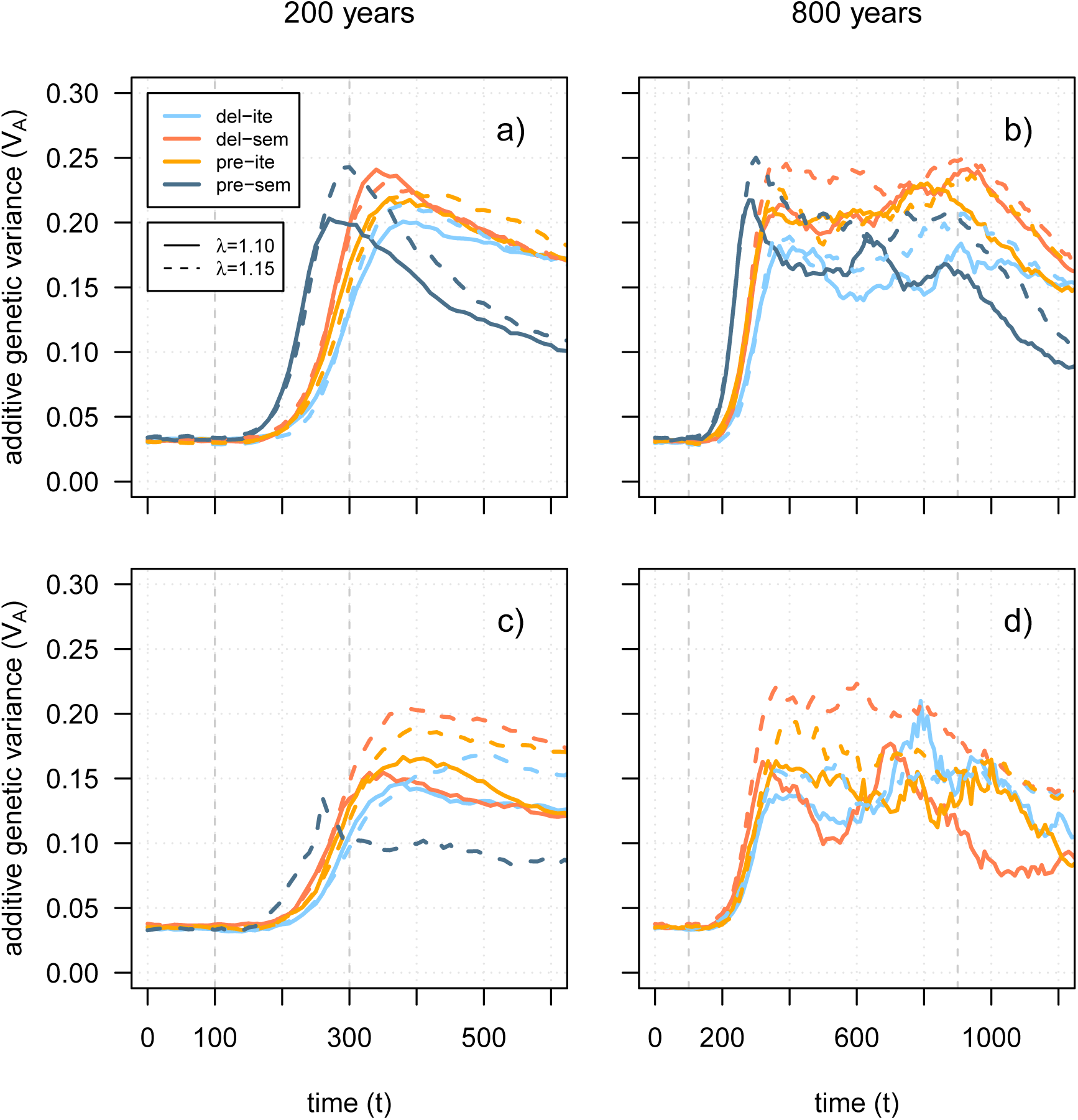
evolution of *G*_2_ with 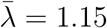. Simulation results for the evolution of additive genetic variance during environmental change are shown for the four life histories, the two different duration of environmental change (200 years - a,c; 800 years - b,d) and two degrees of interannual environmental variability (*ϵ* = 0.2 - a,b; *ϵ* = 1.0 - c,d). The simulation results were averaged over all of the 100 replicates that did not go extinct.

**Figure S11:**
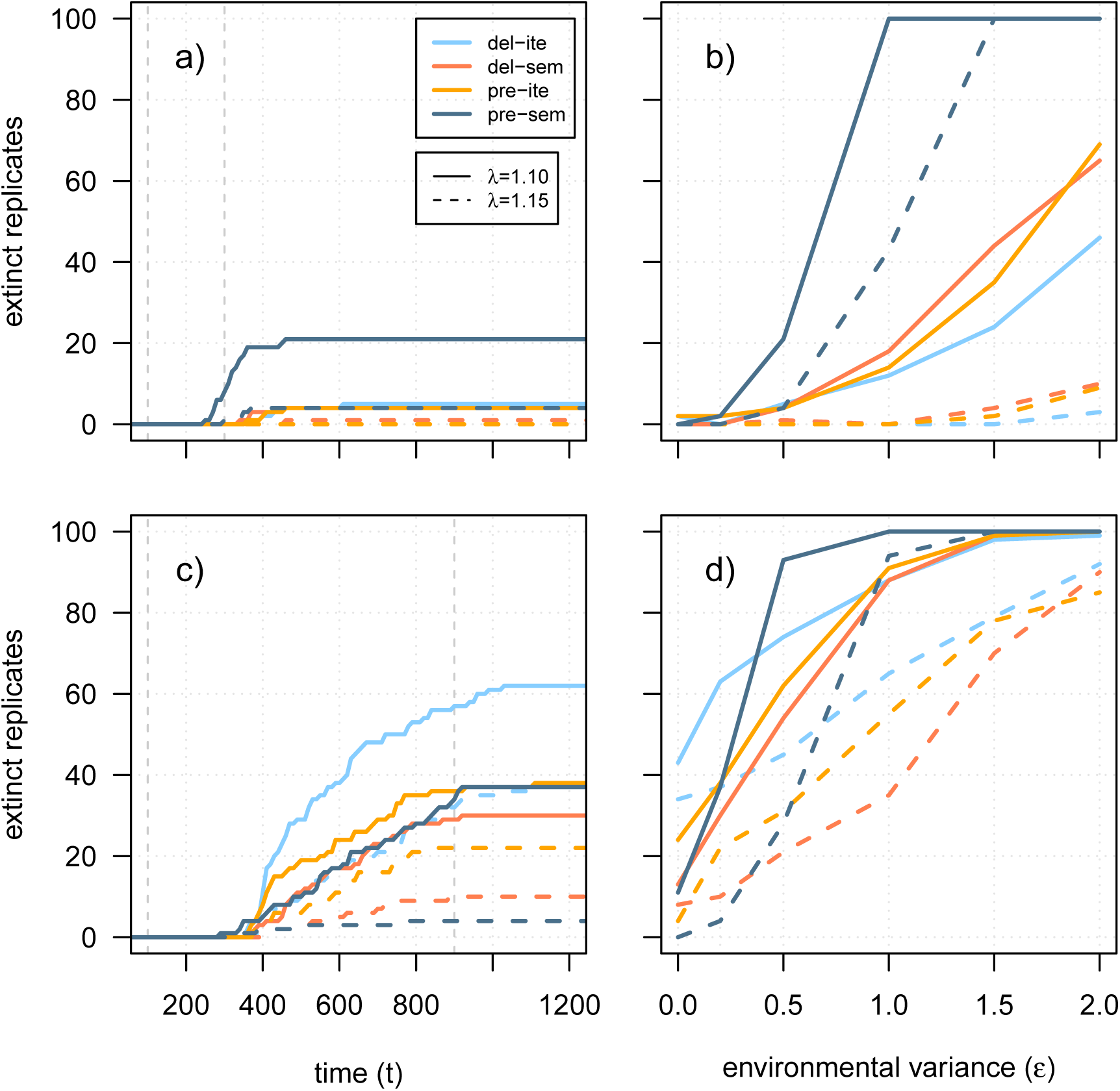
extinction dynamics with 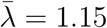. These graphs show the same simulation results on the number of extinct replicates as figure 6, but illustrate additionally the results for the life-history standardization of 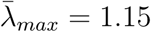. These graphs show for each of the four life histories the number of extinct replicates over time *t* (a,c) and the total number of extinct replicates depending on environmental variability *ϵ* (b,d). Extinction dynamics are shown for two different scenarios with environmental change either lasting for 200 years (a,b) or for 800 years (c,d). The gray dashed lines indicate the start of environmental change (*t* = 100) and its end (*t* = 300 for a,b; *t* = 900 for c,d). In both scenarios, the rate of environmental change per year was set to 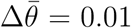. The temporal dynamics are shown for an environmental variability of *ϵ* = 0.5 (a) and *ϵ* = 0.2 (c).

**Figure S12:**
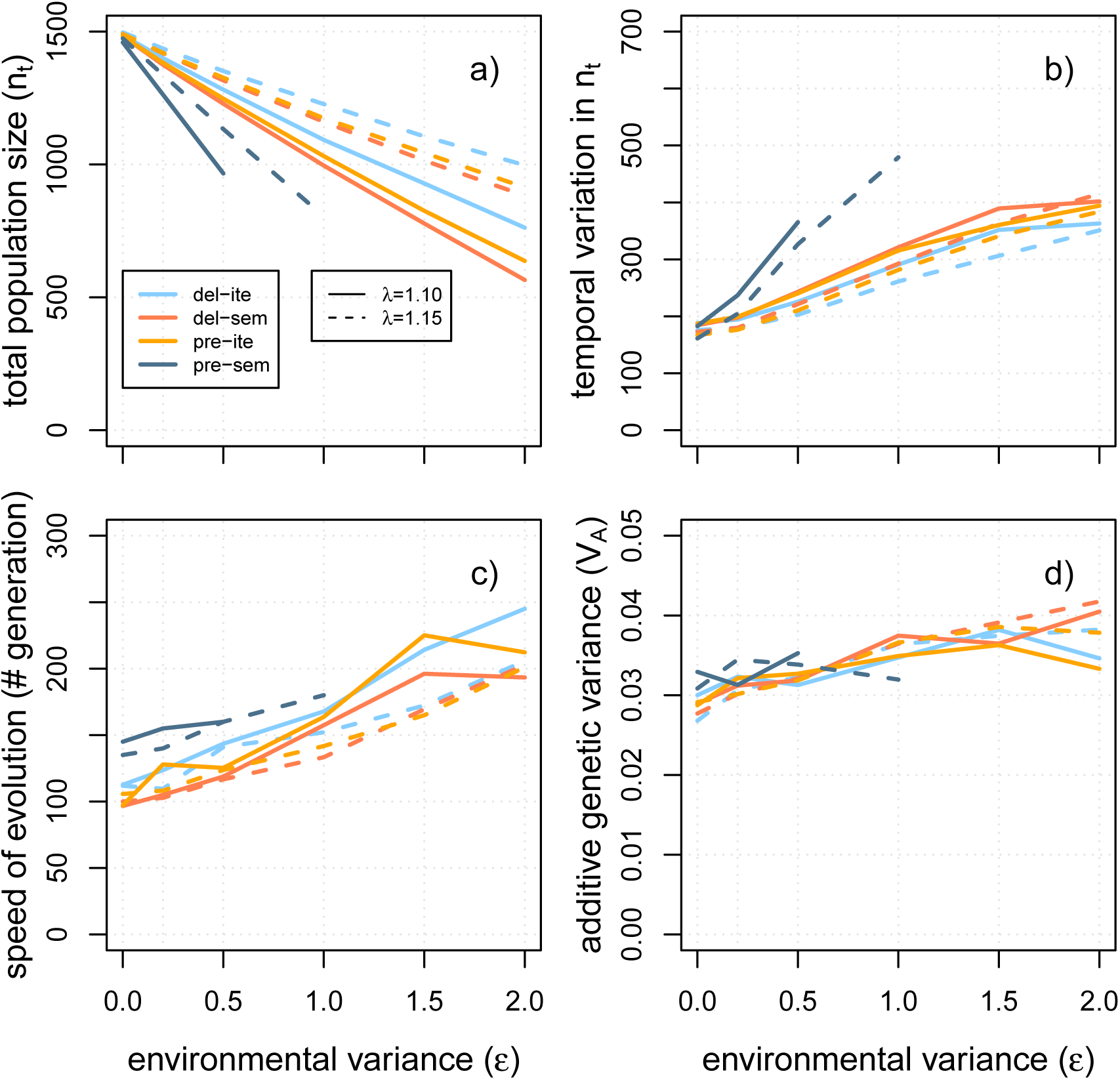
evolvability per generation. These graphs are identical to Fig. 5, except that the speed of evolution (c) is quantified as the number of generations (not years!) to reach 90% of the novel phenotypic optimum after climate change. For this purpose, the number of years to reach the threshold was divided by the average generation time of each life-history strategy at stable conditions from table S1. Please not that the realized generation time in the course of environmental change might differ from these estimates that have been calculated at equilibrium.

**Figure S13:**
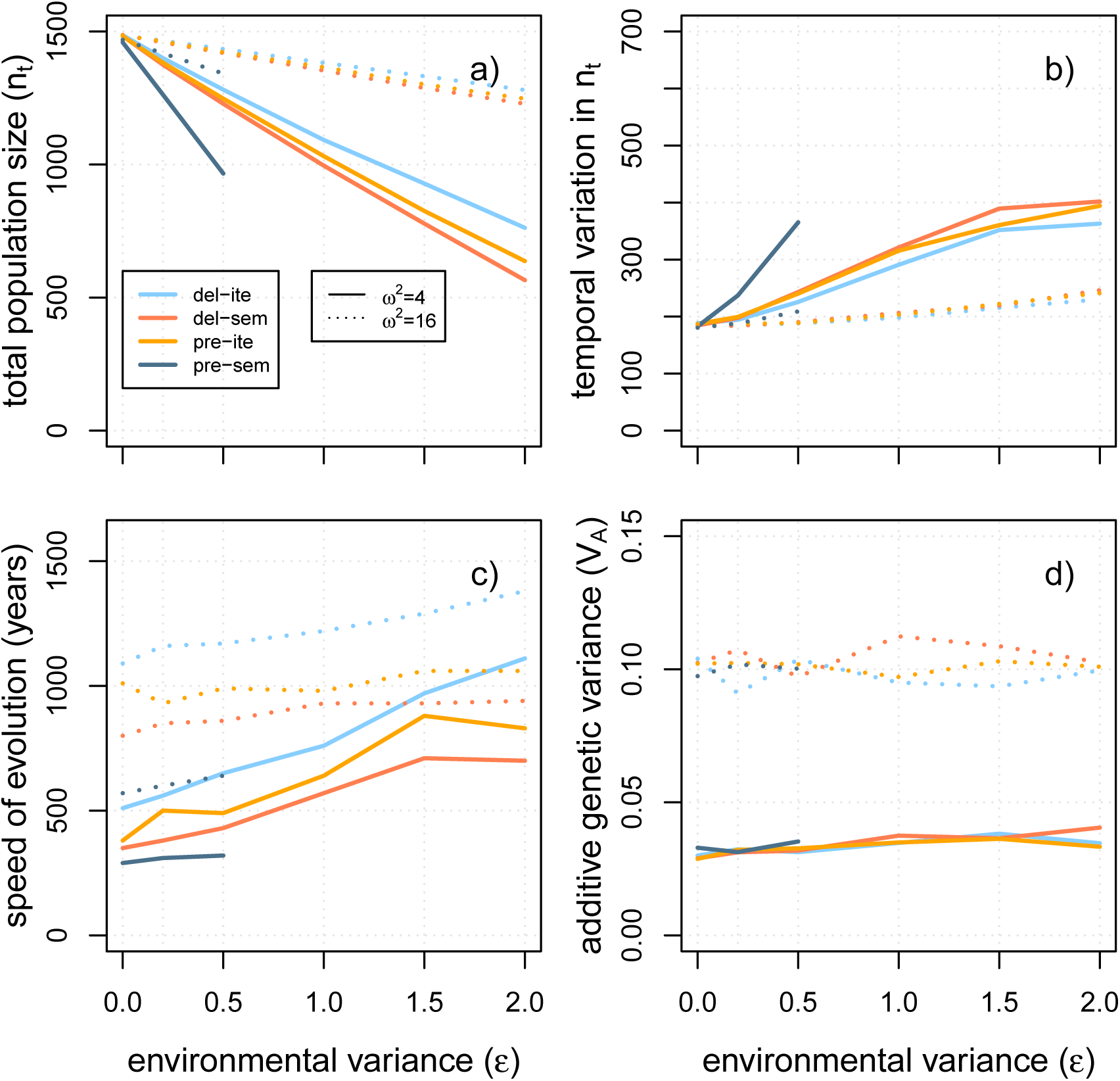
robustness and evolvability with *ω*^2^ = 16. The simulation results for strong selection (*ω*^2^ = 4) are plotted together with the results for weak selection (*ω*^2^ = 16). Weaker selection strength leads to higher population sizes after burn-in (a), lower stochasticity in total population sizes over time (b), a slower speed of evolution (c), and higher levels of additive genetic variance in the juvenile trait (d). However, the rank order of life-history strategies with regard to robustness and evolvability is preserved.

**Figure S14:**
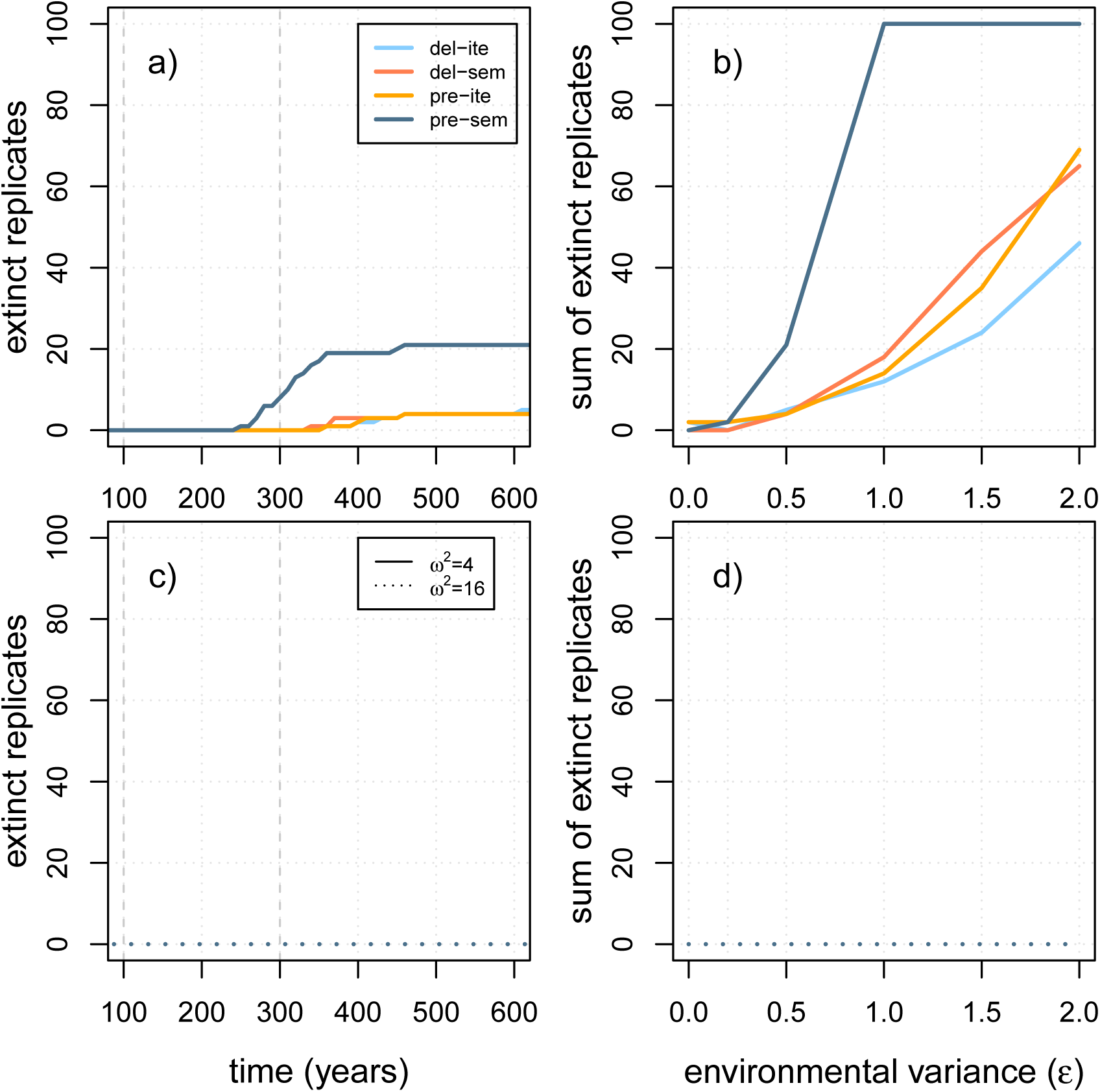
extinction dynamics with *ω*^2^ = 16. The effects of a stronger (*ω*^2^ = 4, a-b) versus weaker (*ω*^2^ = 16, c-d) selection on extinction probabilities is shown for the environmental change scenario over 200 years, similar to Fig. 6.

**Figure S15:**
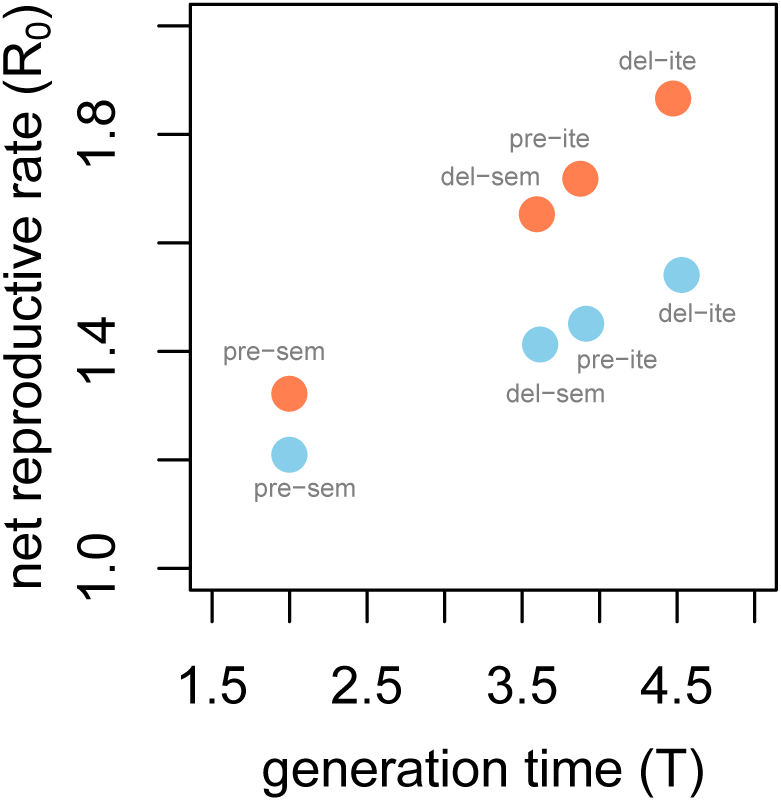
Simulated life histories along fast-slow continuum and reproductive strategy. The generation time (*T*) of each simulated life history is plotted against its net reproductive rate (*R*_0_) from table S1. While the blue circles illustrate life histories standardized for 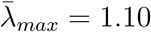, orange dots represent life histories with 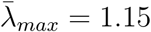. While the x-axis could be interpreted to represent the fast-slow continuum, the y-axis might rather illustrate the reproductive strategy axis as described in meta-analyses (Salguero-Gómez et al., 2017). Please note that both quantities (*T* and *R*_0_) were computed in absence of density regulation.

